# HDAC10 blockade upregulates SPARC expression thereby repressing melanoma cell growth and BRAF inhibitor resistance

**DOI:** 10.1101/2023.12.05.570182

**Authors:** Hongbo Ling, Yixuan Li, Changmin Peng, Shengyu Yang, Edward Seto

**Author notes:** These authors contributed equally (Hongbo Ling, Yixuan Li, Changmin Peng).

## Abstract

Secreted Protein Acidic and Rich in Cysteine (SPARC), a highly conserved secreted glycoprotein, is crucial for various bioprocesses. Here we demonstrate that histone deacetylase 10 (HDAC10) is a key regulator of *SPARC* expression. HDAC10 depletion or inhibition upregulates, while overexpression of HDAC10 downregulates, SPARC expression. Mechanistically, HDAC10 coordinates with histone acetyltransferase p300 to modulate the acetylation state of histone H3 lysine 27 (H3K27ac) at *SPARC* regulatory elements and the recruitment of bromodomain-containing protein 4 (BRD4) to these regions, thereby tuning *SPARC* transcription. HDAC10 depletion and resultant SPARC upregulation repress melanoma cell growth, primarily by induction of autophagy via activation of AMPK signaling. Moreover, SPARC upregulation due to HDAC10 depletion partly accounts for the resensitivity of resistant cells to a BRAF inhibitor. Our work reveals the role of HDAC10 in gene regulation through epigenetic modification and suggests a potential therapeutic strategy for melanoma or other cancers by targeting HDAC10 and SPARC.

**Highlights:**

- HDAC10 is the primary HDAC member that tightly controls SPARC expression.
- HDAC10 coordinates with p300 in modulating the H3K27ac state at *SPARC* regulatory elements and the recruitment of BRD4 to these regions.
- HDAC10 depletion and resultant SPARC upregulation inhibit melanoma cell growth by inducing autophagy via activation of AMPK signaling.
- SPARC upregulation as a result of HDAC10 depletion resensitizes resistant cells to BRAF inhibitors.

## INTRODUCTION

Secreted Protein Acidic and Rich in Cysteine (SPARC), also known as osteonectin (ON) or basement-membrane protein 40 (BM-40), is a highly conserved glycoprotein present across invertebrates and vertebrates. First reported in bone tissues, SPARC possesses the ability to interact with collagen fibrils and hydroxyapatite at discrete locations, underscoring its pivotal role in the process of bone mineralization (1). Later studies identified the presence of SPARC in endothelial cells and a diverse array of other tissues (2, 3), and characterized it as an extracellular matrix (ECM)-associated protein that is vital for tissue remodeling (4), morphogenesis (5, 6), and angiogenesis (7, 8). Over the years, the comprehensive study of SPARC has revealed that it is an integral regulator in a myriad of physiological and pathological processes, including tumorigenesis (9–13).

Depending on the specific cell types and animal models under study, SPARC can manifest contrasting roles in cancer (13). This can be attributed to variations in its expression, secretion, and wide-ranging biological functions. Produced and secreted by various cell types, ranging from cancer cells to stromal and immune cells, SPARC influences the tumor microenvironment in several ways. For example, SPARC plays instrumental roles in the deposition, assembly, and remodeling of ECM, encompassing processes like collagen transformation and matrix protein secretion (14–16). Further, by interactions with soluble factors and consequent alteration in signaling pathways, SPARC impacts proliferation and migration in cancer cells (7, 17–19). In addition, SPARC can alter the tumor microenvironment by dampening immune cell infiltration (20, 21). While it is most often encountered as a secreted glycoprotein, SPARC is also localized on the cell surface and within the intracellular compartment in several cell types. Within the cell, particularly when associated with membranes, SPARC plays a role in pivotal cellular signaling pathways, impacting cell proliferation, differentiation, and survival (22, 23). Its interactions with cytoskeletal elements, such as actin and tubulin, underline its role in the regulation of migration and structural organization (24–26).

Despite SPARC’s multifaceted roles across various cancer types, it’s undeniable that modifying its expression and activity can significantly alter tumor and cellular characteristics. For instance, SPARC inhibits proliferation in ovarian cancer (27, 28), breast cancer (29), and medulloblastoma (30). Furthermore, it mitigates peritoneal carcinomatosis and metastasis in these cancers (31). Consistently, exogenous SPARC administration has demonstrated a suppression of growth in a spectrum of cancers including pancreatic cancer (32), colorectal cancer (33), neuroblastomas (34), and leukemia (35). SPARC has also been implicated in chemoresistance. In the realm of colorectal cancer, SPARC enhances the chemosensitivity of resistant cells either paired with chemotherapy agents (33) or combined with vitamin D (36).

Similarly, high SPARC expression is associated with better outcomes of non-small cell lung cancer (NSCLC) patients treated with cisplatin (37). Accordantly, highly expressed and exogenous SPARC displayed an increased susceptibility of melanomas to the growth-inhibitory impact of drugs like cisplatin (38). These collective findings bolster the idea of targeting SPARC therapeutically in various malignancies, and a deeper exploration into the molecular mechanisms that regulate SPARC expression in cancers is imperative.

Histone deacetylases (HDACs) are enzymes that catalyze the removal of acetyl groups from ε-N-acetyl lysine residues on histone and non-histone proteins (39, 40). In humans, 18 HDAC enzymes have been identified and are grouped into four classes based on their similarity to yeast homologs. Class I consists of HDACs 1‒3, and 8, which are highly homologous to the yeast HDAC Rpd3. Class II comprises HDACs 4‒7, 9, and 10, with their deacetylase domains closely related to that of yeast Hda1. Class III includes seven members of the sirtuin proteins (SIRT1‒7) that are related to the silent information regulator 2 (Sir2). HDAC11 is the sole member of class IV. While classes I, II, and IV are zinc-dependent enzymes showing sequence similarity to each other, they lack homology with Sir2-related proteins that require NAD^+^. The reversible acetylation and deacetylation of histones play a pivotal role in gene expression regulation. Generally speaking, when histone acetyltransferases (HATs) acetylate histones, the resultant neutralization of the lysine residues diminishes histone-DNA affinity, leading to chromatin loosening and subsequent gene transcription. In opposition, deacetylation by HDACs enhances histone-DNA interaction, compacting the DNA and repressing transcription (41).

HDACs and HATs also modulate the acetylation state of numerous non-histone proteins, which impacts the protein’s structure, function, or interactions, often altering gene transcription (39, 42). In addition, HDACs might not change the acetylation state of histone and non-histone protein partners. Instead, they may regulate gene transcription by directly interacting with and, in turn, affecting the function of transcription (co)factors (43). In essence, while HDACs are known for their diverse gene regulatory roles and implications in both cancer and non-cancerous diseases, their specific influence on SPARC expression, particularly in the cancer landscape, remains unknown.

In this study, we demonstrate that HDAC10 is pivotal in orchestrating SPARC expression. HDAC10 blockade resulted in an elevated H3K27ac level and augmented recruitment of bromodomain-containing protein 4 (BRD4) at *SPARC* regulatory elements, thereby promoting *SPARC* transcription. Furthermore, our work shows that HDAC10 depletion and resultant SPARC upregulation led to repressive melanoma cell growth and increased vulnerability to BRAF inhibition in resistant cells. These findings uncover a novel mechanism by which HDAC10 regulates gene expression through histone modification and suggest that targeting HDAC10 and SPARC may hold therapeutic potential in combating melanoma and other malignancies.

## MATERIALS AND METHODS

### Cell Lines

The following cell lines were used in this study: HEK293T (ATCC, CRL-3216, RID:CVCL_0063), A375 (ATCC, CRL-1619, RRID:CVCL_0132), SK-MEL-2 (ATCC, HTB-68, RRID:CVCL_0069), SK-MEL-5 (ATCC, HTB-70, RRID: CVCL_0527), SK-MEL-28 (ATCC, HTB-72, RRID:CVCL_0526), HMEL-BRAF^V600E^ (RRID:CVCL_0132), UACC257 (RRID:CVCL_1779), WM983A (RRID:CVCL_6808), WM3918 (RRID:CVCL_0526), 1205Lu (RRID: CVCL_5239), WM793 (RRID: CVCL_8787), A549 (ATCC, CCL-185, RRID:CVCL_0023), and H1299 (ATCC, CRL-5803, RRID:CVCL_0060).

All these cells were grown in Dulbecco’s Modified Eagle Medium (DMEM) supplemented with 10% fetal bovine serum (FBS) and 1% penicillin-streptomycin at 37°C in a humidified atmosphere of 5% CO2.

### Antibodies

The antibodies used in this study are: anti-HDAC10 (Sigma-Aldrich, #H3413, RRID:AB_261940), anti-FLAG epitope (M2; Sigma-Aldrich, #F1804, RRID:AB_262044), anti-alpha-tubulin (12G10; Developmental Studies Hybridoma Bank; RRID:AB_2315509), anti-beta-actin (Proteintech, #66009-1-IG, RRID:AB_2687938), anti-V5 epitope (Thermo Fisher Scientific, #MA5-15253, RRID:AB_10977225), anti-SPARC (Cell Signaling Technology, #5420, RRID:AB_10692794), anti-acetyl-histone H3 (Lys27) (Cell Signaling Technology, #4353, RRID:AB_10545273), anti-Histone H3 (Cell Signaling Technology, #9715, RRID:AB_331563), anti-BRD4 (Cell Signaling Technology, #13440, RRID:AB_2687578), anti-phospho-AMPK (T172) (Cell Signaling Technology, #2531, RRID:AB_330330), anti-AMPK (Cell Signaling Technology, #2532, RRID:AB_330331), anti-phospho-acetyl-CoA carboxylase (Ser79) (Cell Signaling Technology, #3661, RRID:AB_330337), anti-acetyl-CoA carboxylase (Cell Signaling Technology, #3662, RRID:AB_2219400), anti-phospho-Tuberin/TSC2 (Ser1387) (Cell Signaling Technology, #5584, RRID:AB_10698883), anti-Tuberin/TSC2 (Cell Signaling Technology, #4308, RRID:AB_10547134), anti-phospho-p70 S6 Kinase (Thr389) (Cell Signaling Technology, #9234, RRID:AB_2269803), anti-p70 S6 Kinase (Cell Signaling Technology, #2708, RRID:AB_390722), anti-phospho-ERK1/2 (Thr202/Tyr204) (Cell Signaling Technology, #9101, RRID:AB_331646), anti-ERK1/2 (Cell Signaling Technology, #9102, RRID:AB_330744), anti-phospho-AKT (Ser473) (Cell Signaling Technology, #4060, RRID:AB_2315049), anti-phospho-AKT (Ser473) (Cell Signaling Technology, #9272, RRID:AB_329827), anti-phospho-c-JUN (Ser63) (Cell Signaling Technology, #2361, RRID:AB_490908), anti-c-JUN (Cell Signaling Technology, #9165, RRID:AB_2130165), anti-phospho-SMAD2 (Ser465/Ser467) (Cell Signaling Technology, #18338, RRID:AB_2798798), anti-SMAD2 (Cell Signaling Technology, #5339, RRID:AB_10626777), anti-phospho-SMAD3 (Ser423/425) (Cell Signaling Technology, #9520, RRID:AB_2193207), anti-SMAD3 (Cell Signaling Technology, #9523, RRID:AB_2193182), anti-Beclin-1 (Cell Signaling Technology, #3495, RRID:AB_1903911), anti-p62 (Cell Signaling Technology, #5114, RRID:AB_10624872), anti-LC3A/B (Cell Signaling Technology, #12741, RRID:AB_2617131), anti-PARP (Cell Signaling Technology, #9542, RRID:AB_2160739), anti-cyclin D1 (Santa Cruz Biotechnology, #sc-753, RRID:AB_2070433), anti-cyclin E (Santa Cruz Biotechnology, #sc-198, RRID:AB_631346),anti-cyclin A (Santa Cruz Biotechnology, #sc-751, RRID:AB_631329), anti-cyclin B1 (Santa Cruz Biotechnology, #sc-752, RRID:AB_2072134), anti-p21 (Santa Cruz Biotechnology, #sc-397, RRID:AB_632126), anti-p27 (Santa Cruz Biotechnology, #sc-527, RRID:AB_632131), anti-p300 (Santa Cruz Biotechnology, #sc-585, RRID:AB_2231120), anti-vinculin (Santa Cruz Biotechnology, #sc-73614, RRID:AB_1131294), and anti-Caspase 3 (BD Biosciences, #610322, RRID:AB_397713).

### Chemical Reagents

Trichostatin A (#T8552), Bufexamac (#B0760), Tubacin (#SML0065), MS-275 (#EPS002), and Sodium butyrate (#B5887) were obtained from Sigma-Aldrich. Vemurafenib (PLX4032, #RG7204) was purchased from Selleck Chemicals. A-485 (#24119) was from Cayman Chemical Company.

### Plasmids, transfection, and lentiviral transduction

The lentiviral vector pLenti-CMV-Neo-DEST (Addgene, #17392, RRID:Addgene_17392) was a gift from Eric Campeau & Paul Kaufman (108).The pLenti-CMV-Neo-DEST-HDAC10 was described previously (57). The plasmid lentiCRISPR v2 (Addgene, # 52961, RRID:Addgene_52961) was a gift from Feng Zhang (109). HDAC10 CRISPR/Cas9 knockout plasmid was generated by inserting gRNA sequences targeting 5’-GACGCTCGATCTCGCACTCGGGG-3’ into lentiCRISPR v2. The pLKO.1-based lentiviral plasmids for delivering shRNAs targeting human HDAC1‒11, SPARC, p300 and BRD4 were purchased from Sigma-Aldrich. The plasmid pLKO.1 GFP shRNA (Addgene, #30323, RRID:Addgene_30323) was a gift from David Sabatini (110).The targeting sequences are described in Supplemental Table S1. The plasmids pcDNA-dCas9-p300 Core (Addgene, #61357, RRID:Addgene_61357) and pcDNA-dCas9-p300 Core (D1399Y) (Addgene, #61358, RRID:Addgene_61358) were gifts from Charles Gersbach (111). The plasmid pCDNA-dCas9-HDAC10 was generated by replacing the p300 core encoding sequence of plasmid pcDNA-dCas9-p300 Core with HDAC10 cDNA. The plasmid pLX304-SPARC (#HsCD00444770) was purchased from DNASU.

Lentivirus was prepared by transfecting HEK293T cells with the lentiviral plasmid together with packaging vectors. Lentivirus-transduced cells were selected for neomycin (G418) or puromycin resistance.

### In vivo tumor growth

For assessment of the effect of HDAC10 on melanoma growth, control and HDAC10 knockdown A375 cells (1×10^6^ in 100 μl PBS) were subcutaneously injected into the flanks of NSG mice at the age of 6–8 weeks (Jackson Laboratory, RRID:IMSR_JAX:005557). The mice were sacrificed 3 weeks after injection and the tumor weight was measured. For the investigation of lung and liver colonization by melanoma cells, cells were injection into NSG mice via tail vein. Mice were euthanized 4–5 weeks post-injection. The influence of HDAC10 on lung and liver colonization of melanoma cells was assessed by measuring body weight, lung weight, calculating the ratio of lung weight to body weight, and counting the number of surface tumors. This was complemented by a subsequent histopathological analysis.

Ethical Compliance: All animal experiments were conducted in accordance with the ethical guidelines and approved by the Institutional Animal Care and Use Committee of The George Washington University.

### Real-time quantitative PCR (RT-qPCR)

Total RNA was extracted from cells with the mirVana miRNA isolation kit (Ambion, #AM1560) or TRIzol Reagent (Thermo Fisher Scientific, #15596018), and was reverse transcribed into cDNA using the qScript cDNA synthesis kit (Quanta Biosciences, #95047). Relative quantitation of mRNAs was carried out via SYBR green-based quantitative PCR using a Quantstudio 3 Real-time PCR system (Applied Biosystems). PCR results were analyzed using the 2^-(ΔΔCT)^ method.

The PCR primers sequences are shown in Supplemental Table S1.

### Colony formation assay

Cells were seeded on 3.5-cm dishes or 6-well plates. After appropriate periods, cells were washed with PBS and stained with 1% crystal violet for 30 min at room temperature. The resultant colonies were subsequently counted, imaged, and the relative growth was quantified using Image J software (RRID:SCR_003070).

### Spheroid growth

Three-dimensional spheroids were done as described previously (55). Briefly, 96-well tissue culture plates were pre-coated with 75 μl of 1% agarose in PBS. Nearly confluent cells were trypsinized and seeded at 10^4^ cells/well in 150 μl of DMEM to obtain a single homotypic spheroid per well. Every 3 days, 75 μl of supernatant was delicately removed from each well and replenished with fresh medium. On day 6, the spheroid size was evaluated using an inverted microscope. Spheroid volume was determined using the equation *V*=(*LW*^2^) × 0.5, where *L* is length and *W* is width of the spheroid.

### Immunoblotting

Protein samples (cell lysates or media) were resolved by SDS-PAGE, and electrotransferred onto a nitrocellulose membrane. The blots were incubated in blocking buffer [5% (w/v) nonfat dry milk in TBS + 0.05% Tween 20 (TBS-T)] for 1 h, and incubated with primary antibodies for 16 h at 4°C, followed by incubation with horseradish peroxidase (HRP)-linked goat anti-rabbit IgG (Cell Signaling Technology, #7074, RRID:AB_2099233) or HRP-linked goat anti-mouse IgG (Cell Signaling Technology, #7076, RRID:AB_330924) for 1 h at room temperature. The antibody-antigen complex was visualized by ECL chemiluminescence.

### Indirect immunofluorescence

Cells were fixed and permeabilized. Cells were blocked by 10% normal goat serum (Sigma-Aldrich, #G9023) in PBS for 1 h, then were incubated with primary antibodies for 1 h and with either Alexa Fluor^TM^ 594-conjugated goat anti-mouse IgG (H+L; Invitrogen, #A11005, RRID:AB_2534073) or Alexa Fluor^TM^ 488-conjugated goat anti-rabbit IgG (H+L; Invitrogen, #A11008, RRID:AB_143165) for 1 h. All immunostaining experiments in this study were accompanied with DAPI staining. The BrdU-incorporation assay was performed using the BrdU Labeling and Detection Kit I (Roche, #11296736001, RRID:AB_2814711) according to the manufacturer’s instruction.

### Chromatin immunoprecipitation (ChIP) assay

A ChIP assay was performed essentially as described previously (57). Briefly, cells were incubated for 10 min in 1% formaldehyde, followed by addition of 0.125 M glycine and sonication to generate DNA fragments of 200–1,000 bp. Chromatin from about 1.5×10^6^ cells was incubated overnight at 4°C with 2–5 μg antibodies to FLAG epitope (M2; Sigma-Aldrich, #F1804, RRID:AB_262044), BRD4 (Cell Signaling Technology, #13440, RRID:AB_2687578), or acetyl-histone H3 (Lys27) (Cell Signaling Technology, #4353, RRID:AB_10545273). Normal mouse IgG (Santa Cruz Biotechnology, #sc-2025, RRID:AB_737182) and normal rabbit IgG (Millipore, #12-370, RRID:AB_145841) were used as controls. Samples were incubated with preblocked protein A/G Plus beads (Santa Cruz Biotechnology, #2003, RRID:AB_10201400) for 2 h at 4°C. DNA was purified and then amplified by PCR using the primers shown in Supplemental Table S1.

### Statistical analysis

All statistical analyses were carried out using GraphPad Prism 8(RRID:SCR_002798) or Microsoft Excel. The normality of continuous variables was analyzed by Shapiro-Wilk test. For normally distributed data, comparisons were made using the un-pooled two-sample *t*-test.

Differences were considered statistically significant when the *P*-value was less than 0.05.

## RESULTS

### HDAC10 represses SPARC expression in melanoma cells

To explore the potential relationship of HDACs and SPARC, we knocked down individual classical HDACs (HDAC1‒11) in A375 melanoma cells using lentivirus-delivered short hairpin RNA (shRNA) (Figure S1). SPARC mRNA levels were then evaluated. The knockdown of HDAC10 resulted in a marked increase in the level of *SPARC* mRNA, whereas the knockdown of HDAC5 resulted in a modest increase in *SPARC* mRNA. There were no significant SPARC RNA changes from depletion of the other nine HDACs. Thus, HDAC10 is the major HDAC that suppresses SPARC expression in melanoma cells (Figure 1A). Concordantly, knockdown (Figure 1B) or knockout (Figure 1C) of HDAC10 in A375 cells resulted in robustly elevated intracellular and extracellular accumulation of SPARC protein. Similar results were obtained when other melanoma cell lines were depleted of HDAC10 (Figure 1D). In agreement with these findings, treatment with selective HDAC10 (class IIb HDAC) inhibitor Bufexamac (44) led to an increase in the level of SPARC protein in a dose-dependent manner (Figure 1E).

**Figure 1.**
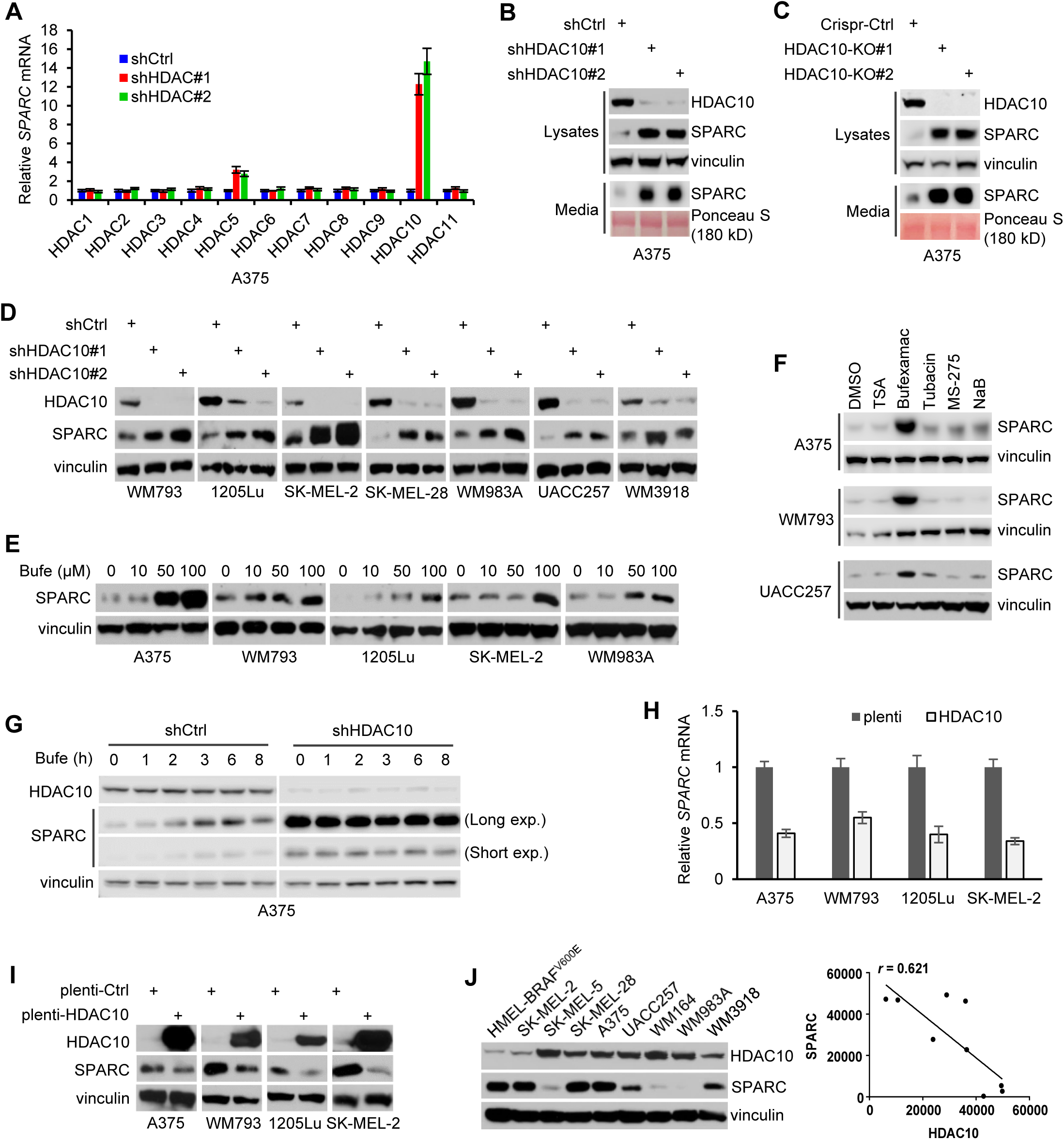
HDAC10 represses SPARC expression in melanoma cells. **(A)** A375 melanoma cells were infected with lentiviruses harboring shRNA targeting individual HDAC (HDACs 1–11). Relative levels of *SPARC* mRNA of the cells were determined by qPCR. **(B)** A375 cells were infected with lentiviruses harboring HDAC10 or control shRNA. Cell lysates were immunoblotted with antibodies to HDAC10, SPARC, and vinculin (loading control). Additionally, culture media proteins were separated, stained with Ponceau S, and immunoblotted for SPARC. **(C)** A375 cells were deleted of HDAC10 by CRISPR/Cas9-mediated knockout. The SPARC expression of the cells was examined by immunoblotting. **(D)** HDAC10 was depleted and SPARC expression was examined across multiple melanoma cell lines. **(E)** Melanoma cells were treated with HDAC10 inhibitor (Bufexamac) or solvent control for 24 h, and SPARC expression was examined. **(F)** Melanoma cells were exposed to different types of HDAC inhibitors (100 ng/ml TSA, 0.1 mM Bufexamac, 2 µM tubacin, 1 µM MS-275, 5 mM sodium butyrate) or control for 24 h, then were examined for SPARC expression. **(G)** HDAC10-depleted and control A375 cells were treated with 0.1 mM Bufexamac for 1–8 h, then were examined for SPARC expression. **(H)** HDAC10 was forcedly expressed, and the relative level of *SPARC* mRNA was quantified. **(I)** Lysates from melanoma cells overexpressing HDAC10 or control were immunoblotted for SPARC. **(J)** Immunoblotting (left panel) and correlation (right panel) analyses of HDAC10 and SPARC protein levels in a panel of melanoma cell lines.

Corroboratively, when HDAC10 was depleted, the effect of Bufexamac treatment on SPARC expression was largely blunted (Figure 1G). In contrast to Bufexamac treatment, exposure to either the broad spectrum HDAC inhibitor Trichostatin A (TSA), or the HDAC6-specific inhibitor Tubacin, or the class I HDACs inhibitors MS-275 and sodium butyrate (NaB), none of which efficiently target HDAC10, has little effect on SPARC expression (Figure 1F). Conversely, overexpression of HDAC10 reduced the level of both *SPARC* mRNA (Figure 1H) and protein (Figure 1I). Thus, HDAC10 suppresses SPARC expression in melanoma cells. In support of this notion, the level of endogenous HDAC10 protein is statistically inversely correlated with endogenous SPARC protein expression in a panel of melanoma cell lines (Figure 1J).

### HDAC10 modulates the H3K27ac state at *SPARC* regulatory elements

Consistent with our results that HDAC10 represses SPARC expression in melanoma cells, in an RNA-seq analysis, SPARC is one of the top upregulated genes upon HDAC10 depletion in H1299 lung cancer cells (Figure S2A). Notably, the basal *SPARC* expression varies across distinct cancer and/or cell types, e.g., the relative level of *SPARC* mRNA in melanoma cells (A375 and WM793) is ∼ 400 to 1,000 times higher than in lung cancer cell lines (H1299 and A549) (Figure S2B). HDAC10 depletion consistently led to a significant increase in the expression of *SPARC* mRNA, regardless of their basal levels (Figure S2C). Therefore, the transcriptional repression of *SPARC* by HDAC10 is not cell-type dependent. Alternatively, HDAC10 may target certain machinery that is commonly shared across cell lines, such as histones. Histones are known to possess multiple acetylation sites, including lysine residues on histones H3, H4, H2A, and H2B. Among these acetylation sites, histone H3 lysine 27 acetylation (H3K27ac), primarily catalyzed by p300/CBP histone acetyltransferase(s) (45), is a well-established marker of active gene regulatory elements crucial for the initiation and elongation of gene transcription. Interestingly, the promoter (Pro: –121 to –32) and enhancers (Enh1: 1,045 to 1,115; Enh2: 3,171 to 3,331; Enh3: 9,470 to 9,540) of *SPARC* are putatively acetylated on H3K27 according to the prediction of the ENCODE database (Figure 2A). To understand whether HDAC10 can regulate the H3K27ac state at these loci, we performed chromatin immunoprecipitation (ChIP) assays using an antibody to H3K27ac. Compared to a control region (–4,622 to –4,537), H3K27ac was significantly enriched at multiple *SPARC* promoter and enhancers when HDAC10 was depleted (Figure 2B) or pharmaceutically inhibited (Figure 2C). In contrast, ectopic HDAC10 expression reduced the H3K27ac level at these regions (Figure 2D). Collectively, our results suggest HDAC10 plays a role in modulating the H3K27ac state at *SPARC* regulatory elements.

**Figure 2.**
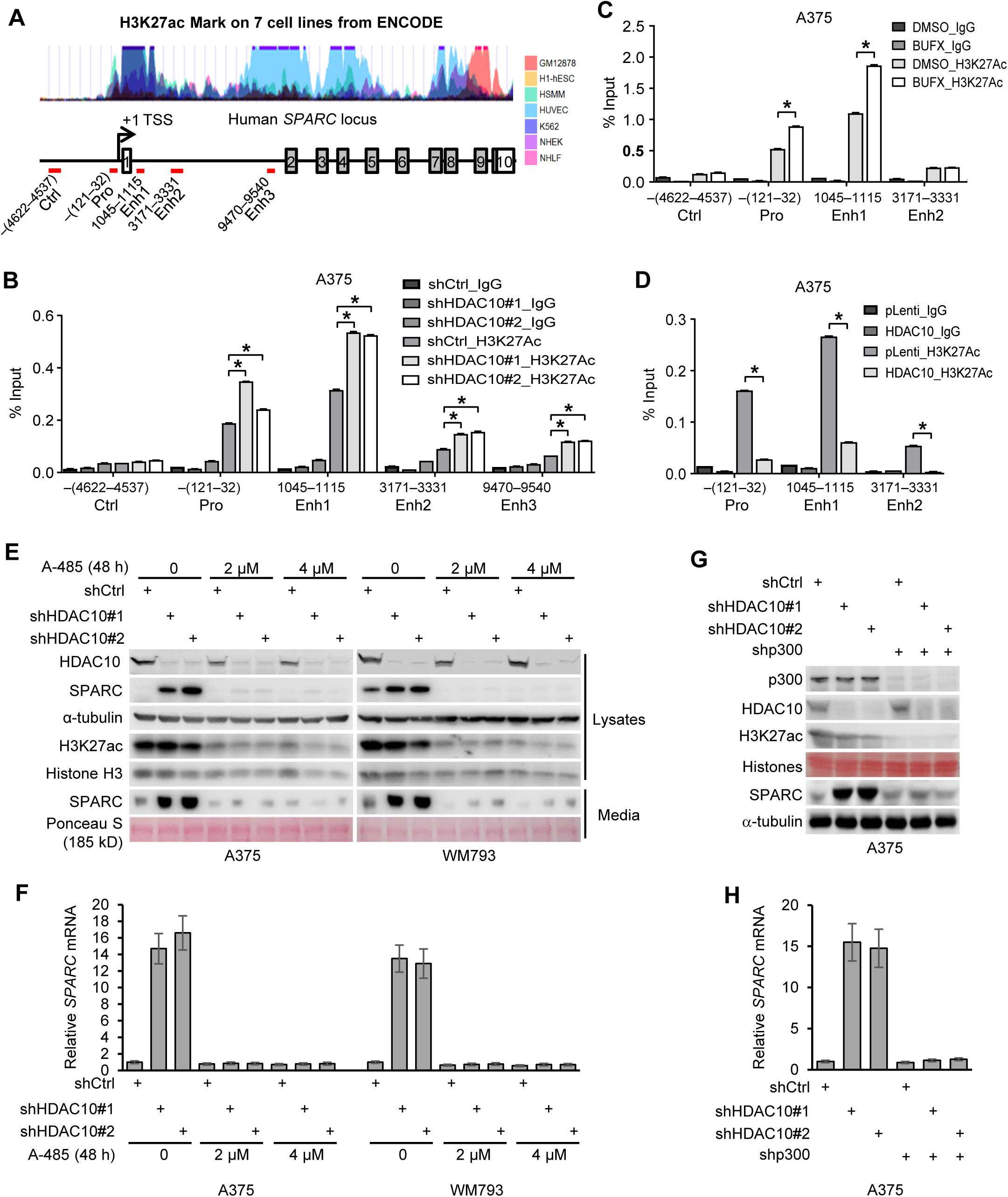
HDAC10 modulates the H3K27ac state at *SPARC* regulatory elements. **(A)** Schematic depiction of the chromatin modification pattern at *SPARC* locus on chromosome 5, adapted from the UCSC Genome Browser database. The +1 position represents the primary transcription initiation site. Red horizontal bars highlight regions amplified by qPCR post-ChIP. **(B)** ChIP analysis of H3K27ac level at *SPARC* locus in HDAC10-depleted and control A375 cells. *, *P* < 0.05. **(C)** A375 cells were overexpressed with HDAC10. The H3K27ac level at *SPARC* locus was assessed by ChIP assays. *, *P* < 0.05. **(D)** Cells were treated with Bufexamac or the control (DMSO). The H3K27ac level at *SPARC* regulatory elements was evaluated by ChIP. *, *P* < 0.05. **(E, F)** HDAC10-depleted and control A375 and WM793 cells were treated with p300/CBP inhibitor A-485 (2 or 4 µM) for 48 h. Cell lysates were analyzed for intracellular levels of HDAC10, SPARC, H3K27ac, and Histone H3. Additionally, secreted SPARC protein from culture media was evaluated (E). SPARC mRNA levels were quantified using qPCR (F). **(G, H)** HDAC10-depleted or control A375 cells were further depleted of p300. The expression of SPARC protein (G) and mRNA (H) was determined.

Given the importance of H3K27ac in gene expression regulation, the observed increase in H3K27ac levels may contribute to the SPARC upregulation by HDAC10 depletion. To corroborate this hypothesis, cells were depleted of HDAC10 and/or treated with A-485, a selective p300/CBP inhibitor (46). The H3K27ac level and the SPARC expression were then examined. Exposure to A-485 blocked the H3K27 acetylation and entirely abolished the SPARC upregulation (Figures 2E, 2F). Consistently, depletion of p300, which significantly reduced the overall level of H3K27ac in cells, attenuated the SPARC upregulation (Figures 2G, 2H); suggesting that HDAC10 regulates *SPARC* transcription, at least partially, by modulating the H3K27ac state through its interplay with p300/CBP.

### Bidirectional regulation of the H3K27ac state and BRD4 accumulation at *SPARC* **regulatory elements by dCas9-HDAC10 and dCas9-p300**

It is important to note that HDAC10 depletion did not significantly alter the overall level of H3K27ac in cells (Figures 2E, 2G). HDAC10 might work in tandem with p300/CBP to fine-tune transcription by modulating local H3K27ac accumulation at *SPARC* locus. To validate this hypothesis, we employed a CRISPR-based approach to selectively tether catalytically inactive Cas9 (dCas9)-fused HDAC10, p300 (core) WT or H1399Y (acetyltransferase inactive mutant) to the putative promoter or enhancers of *SPARC*. The A375 cells, which exhibited high responsiveness of SPARC expression and H3K27ac level to HDAC10 or p300/CBP modulation, were employed for these experiments. Cells were co-transfected with plasmids expressing dCas9-fused proteins and guide RNAs (gRNAs) precisely targeting *SPARC* promoter, enhancers, or control region. The cells transfected with the expression plasmids had a similar expression level of the dCas9-fused proteins (Figure 3A). When co-expressing with gRNAs targeting the promoter, Enh2 or Enh3, the dCas9-p300 and dCas9-HDAC10 led to a significant increase and decrease in *SPARC* mRNA level, respectively. Expression of dCas9-p300-H1399Y did not significantly change *SPARC* mRNA level, reaffirming the instrumental role of the H3K27ac state in modulating *SPARC* expression (Figure 3B). As a control, *HDAC1* expression remained consistent across all co-transfection conditions (Figure 3C), confirming that the alterations in *SPARC* expression can be attributed to the targeted engagement of dCas9-p300 or dCas9-HDAC10 with *SPARC* regulatory elements, as opposed to unintended off-target interactions.

**Figure 3.**
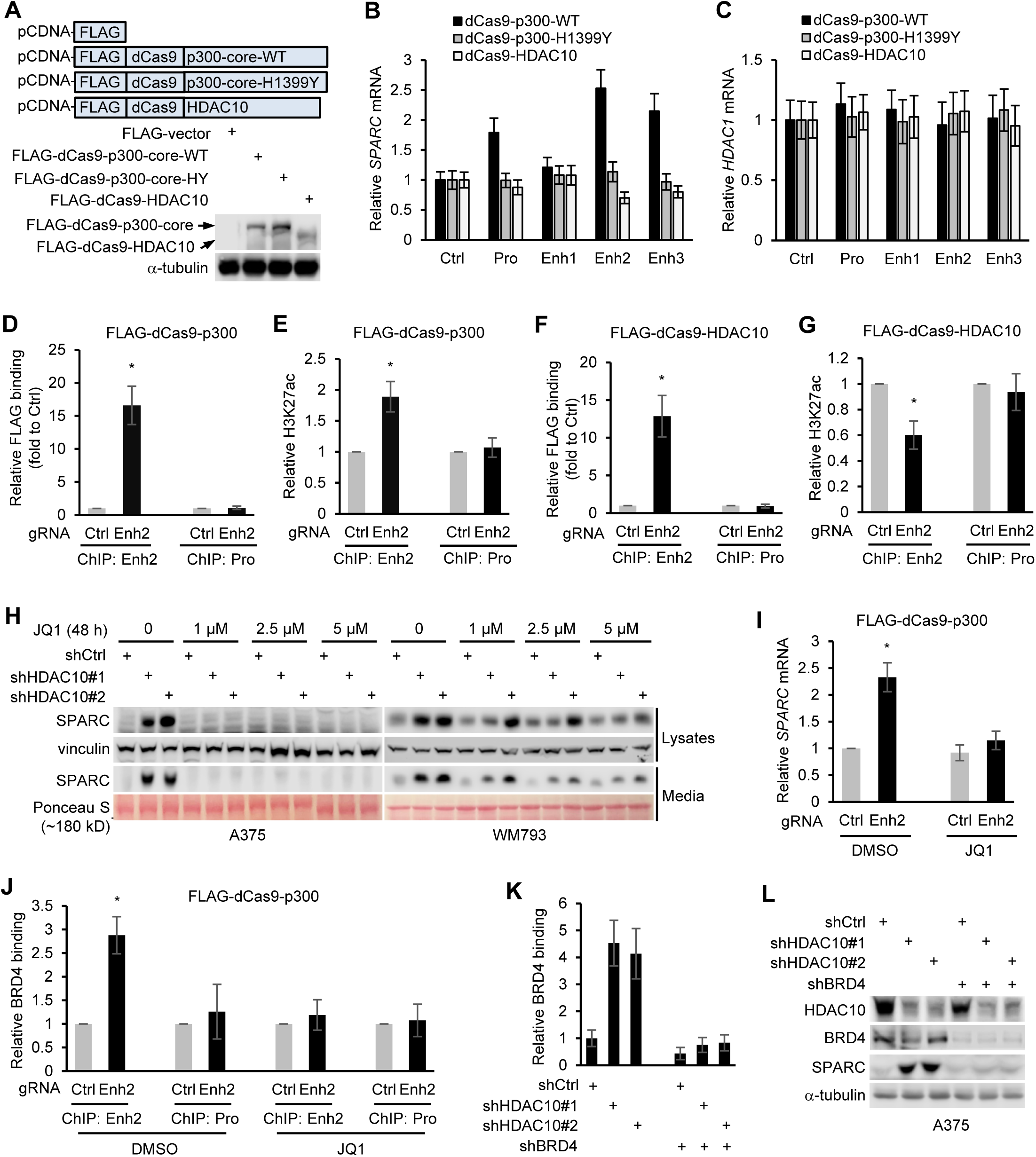
Bidirectional regulation of the H3K27ac state and BRD4 accumulation at *SPARC* regulatory elements by dCas9-HDAC10 and dCas9-p300. **(A)** A375 cells were transfected with plasmids expressing FLAG-tagged dCas9-fused p300 (core WT or H1399Y) or HDAC10. Cell lysates were immunoblotted with antibodies to FLAG epitope and α-tubulin. **(B, C)** Cells were transfected with plasmids expressing dCas9 fused proteins and sgRNAs targeting *SPARC* promoter, enhancer or control regions, respectively. The relative mRNA levels of *SPARC* (B) and *HDAC1* (C) were quantified by qPCR. **(D, E)** Cells were transfected with plasmids expressing FLAG-tagged dCas9-fused p300 and gRNAs targeting Enh2. The relative accumulation of FLAG-tagged dCas9-fused p300 (C) and H3K27ac level (E) at Enh2 and the promoter (serves as control) were determined by ChIP assays with antibodies to FLAG epitope and H3K27ac. *, *P* < 0.05. **(F, G)** Cells were transfected with plasmids expressing FLAG-tagged dCas9-fused HDAC10 and gRNA targeting Enh2. The relative accumulation of FLAG-tagged dCas9-fused HDAC10 (F) and H3K27ac (G) at Enh2 and the promoter were determined by ChIP. *, *P* < 0.05. **(H)** HDAC10-depleted and control cells were treated with JQ1 for 24 h. Cells were washed and maintained in fresh media containing JQ1 for an additional 24 h. Cell lysates and media were collected and blotted for SPARC. **(I, J)** Cells were transfected with plasmids expressing FLAG-tagged dCas9-p300 and gRNAs to Enh2 and control region. Cells were then treated with JQ1 or control. The relative level of *SPARC* mRNA was quantified by qPCR (I). The relative BRD4 binding to Enh2 and control region were evaluated by ChIP with an antibody to BRD4 (J). *, *P* < 0.05. **(K, L)** Cells were depleted of HDAC10 and/or BRD4. The relative BRD4 binding to Enh2 was determined by ChIP (K). Cell lysates were immunoblotted with antibodies to indicated proteins (L).

Three of the four putative regulatory elements played a role in regulating *SPARC* transcription, with Enh2 showing the highest responsiveness to HDAC10 and p300 manipulation (Figure 3B). To investigate whether dCas9-p300 or dCas9-HDAC10 modulates *SPARC* transcription by altering its H3K27ac state, we narrowed our focus to Enh2. Cells co-expressing Enh2 gRNAs along with either dCas9-p300 or dCas9-HDAC10 were subjected to ChIP assays using antibodies to the FLAG epitope and H3K27ac. The recruitment of FLAG-tagged dCas9-fused proteins and the accumulation of H3K27ac at Enh2 and the promoter were then assessed. When co-expressing with Enh2 gRNAs, dCas9-p300 was significantly recruited to Enh2, but not to the promoter (Figure 3D); a significant increase in H3K27ac levels was observed at Enh2, where dCas9-p300 was recruited, but not at the promoter (Figure 3E). In contrast, when dCas9-HDAC10 was recruited to Enh2 (Figure 3F), the H3K27ac level at Enh2 was discernibly decreased. Concurrently, the H3K27ac concentration at the promoter remained relatively stable (Figure 3G). Cumulatively, these observations underline the notion that dCas9-HDAC10 and dCas9-p300 induce a bidirectional influence on the H3K27ac landscape at *SPARC* regulatory elements, thereby steering *SPARC* transcription.

H3K27ac shapes active promoters and enhancers by opening chromatin and recruiting transcription (co)factors to core promoters (47–50). Among these, BRD4 is noteworthy. It recognizes and binds to acetylated histones H3K27ac, and aids in gene transcription by recruiting other (co)factors to the target site and enhancing their functional activity (51–53). Of particular interest is its involvement in SPARC expression regulation; inhibition of BRD4 activity by small molecule inhibitors or RNA interference reduced SPARC expression (54). We wondered if BRD4 plays a role in HDAC10’s regulation of SPARC expression. To address this, cells were depleted of HDAC10 and treated with JQ1, a well-established BRD4 inhibitor that prevents BRD4 from binding to acetylated proteins. Surprisingly, while exposure to JQ1 did not significantly affect the basal expression of SPARC, it markedly blocked the upregulation of SPARC by HDAC10 depletion (Figure 3H). These findings suggest that cells may maintain lower levels of H3K27ac and accumulate less BRD4 at *SPARC* regulatory elements responsible for priming SPARC expression. Hence, BRD4 might not be essential for the priming of SPARC expression. However, upon HDAC10 depletion, the resulting increase in H3K27ac levels may facilitate the recruitment of BRD4 to *SPARC* regulatory elements, such as Enh2, thereby promoting *SPARC* transcription.

To further investigate the potential involvement of H3K27ac in the recruitment of BRD4 to Enh2, cells co-expressing dCas9-p300 and Enh2 gRNAs were treated with JQ1. The *SPARC* expression level and BRD4 occupancy at Enh2 were evaluated. Indeed, the SPARC upregulation by the recruitment of dCas9-p300 to Enh2 requires the activity of BRD4 (Figure 3I). Consistent with the role of BRD4 in enhancer acetylation-dependent transactivation of *SPARC*, the BRD4 occupancy at Enh2 (but not at the promoter) was significantly enriched when dCas9-p300 was recruited to this region (Figure 3J). In line with this, when HDAC10 depletion induced enrichment of BRD4 at Enh2 was attenuated due to BRD4 depletion (Figure 3K), the SPARC upregulation was diminished (Figure 3L); implying the induction of *SPARC* expression is likely due to increased H3K27ac levels and BRD4 occupancy at *SPARC* regulatory elements.

### HDAC10 depletion represses melanoma cell growth by upregulating SPARC expression

Having uncovered HDAC10’s regulation in SPARC expression, we sought to explore its biological implications in cancer cells. SPARC regulates cell growth and proliferation in certain malignancies, including melanoma (55, 56). Accordantly, we observed a marked inhibition in subcutaneous growth (Figure 4A) and a diminished capacity for lung and liver colonization (Figures 4B, 4C) in HDAC10-depleted A375 cells.

**Figure 4.**
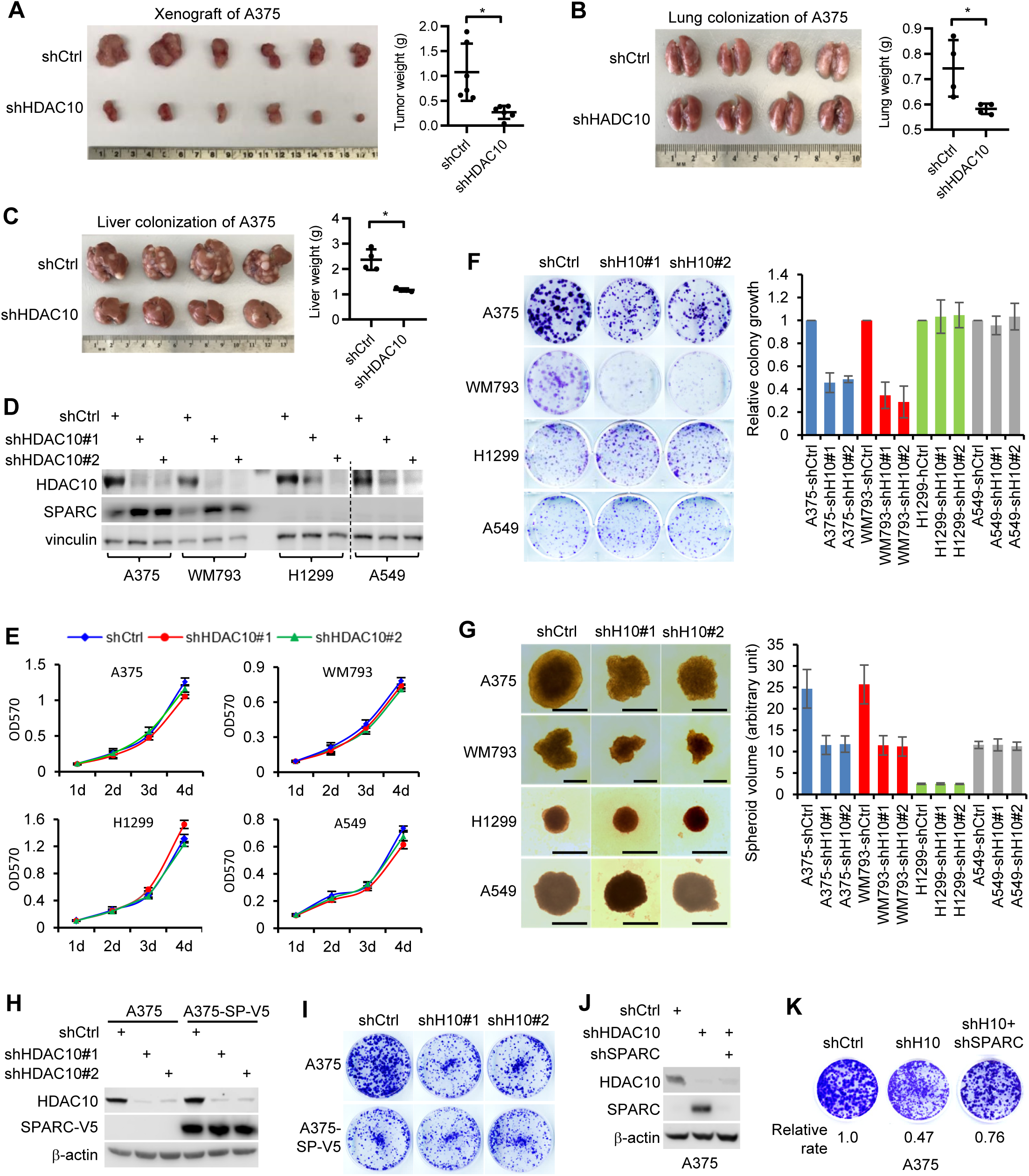
HDAC10 depletion represses melanoma cell growth by upregulating SPARC. **(A)** HDAC10-depleted and control A375 cells (5×10^5^ cells in 100 μl PBS) were subcutaneously injected into NSG mice. Xenograft growth was measured 20 days post-injection. **(B, C)** Lung and liver colonization by HDAC10-depleted and control A375 cells (1×10^6^ cells/injection) at 5 weeks after tail vein injection in NSG mice. The gross morphology of lung (left panel) and the weight of mice lung (right panel) are shown (B). The gross morphology of liver with tumor nodules (left panel) and weight of liver (left panel) are shown (C). **(D)** Following HDAC10 depletion, SPARC protein expression was evaluated. **(E)** Proliferative ability of HDAC10-depleted and control cells was assessed using MTT assay. **(F)** HDAC10-depleted and control cells were evaluated for their colony formation ability. Cells were seeded on 3.5-cm dishes or 6-well plates. For each, 500 (A375, WM793 and H1299) or 1,000 cells (A549) were applied. Cells were stained with crystal violet (left panel) and relative colony growth of cells was quantified by Image J (right panel). **(G)** Spheroid growth of HDAC10-depleted and control melanoma and lung cancer cell lines was assessed. Representative spheroid morphology at day 6 are shown (left panel). Scale bar: 500 μm. Relative spheroid volumes are graphed as means ± SD of at least 10 spheroids (right). **(H)** A375 cells were depleted of HDAC10 and/or overexpressed with V5-tagged SPARC. Cell lysates were immunoblotted for HDAC10 and V5-tagged SPARC. **(I)** Colony growth of A375 cells depleted of HDAC10 and/or overexpressed with SPARC. **(J)** SPARC was further depleted in HDAC10-depleted or control A375 cells, and was confirmed by immunoblotting. **(K)** Colony growth of cells depleted of SPARC and/or HDAC10.

We next dissected the potential involvement of SPARC in HDAC10 depletion induced growth inhibition by MTT, colony formation and spheroid growth assays. It is worth noting that transient depletion of HDAC10 may induce severe cell cycle arrest and growth inhibition via downregulation of cyclin A expression (57). To eliminate this potential confounding factor, we established stable HDAC10-depleted cell lines by long-term culture (∼ 4 weeks), in which the acute cell cycle arrest and growth inhibition are restored. The melanoma (A375 and WM793) and lung cancer (H1299 and A549) cell lines were chosen for this study due to their differential SPARC protein expression levels. While HDAC10 depletion caused a significant SPARC upregulation in melanoma cells, SPARC expressions in lung cancer cells were too low to effectively gauge any change brought about by HDAC10 depletion (Figure 4D). Irrespective of SPARC expression levels, HDAC10 depletion only marginally affected the proliferation of all tested cell lines under 2D culture condition, as determined by the MTT assay (Figure 4E).

However, in melanoma cells characterized by high SPARC expression, HDAC10 depletion significantly repressed colony (Figure 4F) and spheroid (Figure 4G) growth. In contrast, in lung cancer cells where SPARC protein expression is low, HDAC10 depletion had minimal effect on colony or spheroid growth. These results imply that the growth inhibition arising from stable HDAC10 depletion, likely contingent on SPARC, may not profoundly disrupt intrinsic cell cycle progression or induce acute apoptotic cell death. Instead, it is likely that changes associated with cell-cell and/or cell-matrix interactions by HDAC10 depletion play a role in the suppression of colony and spheroid growth. Indeed, SPARC has been shown to regulate a multitude of cellular processes, both via extracellular and intracellular signaling pathways (55, 56).

We then assessed if SPARC is required for the growth suppression by HDAC10 depletion. The A375 cells were ectopically expressed with SPARC and/or depleted of HDAC10 (Figure 4H), and then were examined for growth capacity (Figure 4I). Consistent with previous findings (20, 28, 29), ectopic expression of SPARC resulted in a reduced growth rate and, consequently, overrode the inhibitory effect on colony growth induced by HDAC10 depletion (Figure 4I).

Conversely, when SPARC was depleted (Figure 4J), the colony growth repression by HDAC10 depletion was significantly blunted (Figure 4K). Collectively, HDAC10 depletion suppresses melanoma cell growth, at least in part, by an increase in SPARC expression.

### SPARC upregulation by HDAC10 depletion induces AMPK activation and autophagic cell death

Next, we explored the mechanism underlying the growth repression related to the HDAC10-SPARC axis. SPARC influences multiple signaling pathways, including AMPK, MAPK, and TGFβ/SMAD signaling, among others (17, 19, 58–64). These pathways play critical roles in cell growth, albeit in a cell-context-dependent manner. Therefore, the HDAC10-SPARC axis may regulate the colony and spheroid growth of melanoma cells by interfering with some of these pathways. To test this hypothesis, A375 and WM793 cells were depleted of HDAC10 and were examined for alterations in these signaling pathways by immunoblotting. Upon HDAC10 depletion, the phosphorylation of AMPK, indicative of its activation, was elevated in both cell lines (Figure 5A). Conversely, the activities of other signalings either did not significantly change or showed alterations in only one of the cell lines, making it challenging to attribute the growth inhibition observed in these pathways. Thus, it is likely that AMPK signaling activation, rather than changes in other pathways, primarily contributes to the growth suppression. Consistent with this idea, HDAC10 depletion markedly promoted the phosphorylation of acetyl-CoA carboxylase (ACC), a direct downstream target of AMPK (65), while attenuating the phosphorylation of S6K1, a well-established downstream targets of AMPK, likely through TSC2 phosphorylation (Figure 5B) (66–68). Next, we asked whether the activation of AMPK resulted from the upregulation of SPARC upon HDAC10 depletion. We ectopically expressed SPARC and/or depleted HDAC10 in A375 cells, then examined the phosphorylation of AMPK and its downstream targets. Overexpression of SPARC led to activation of AMPK and its downstream signaling pathways, like HDAC10 depletion (Figure 5C). Conversely, knockdown of SPARC attenuated the activation of AMPK signaling caused by HDAC10 depletion (Figure 5D);suggesting that HDAC10 depletion activates AMPK signaling in a manner dependent on SPARC upregulation.

**Figure 5.**
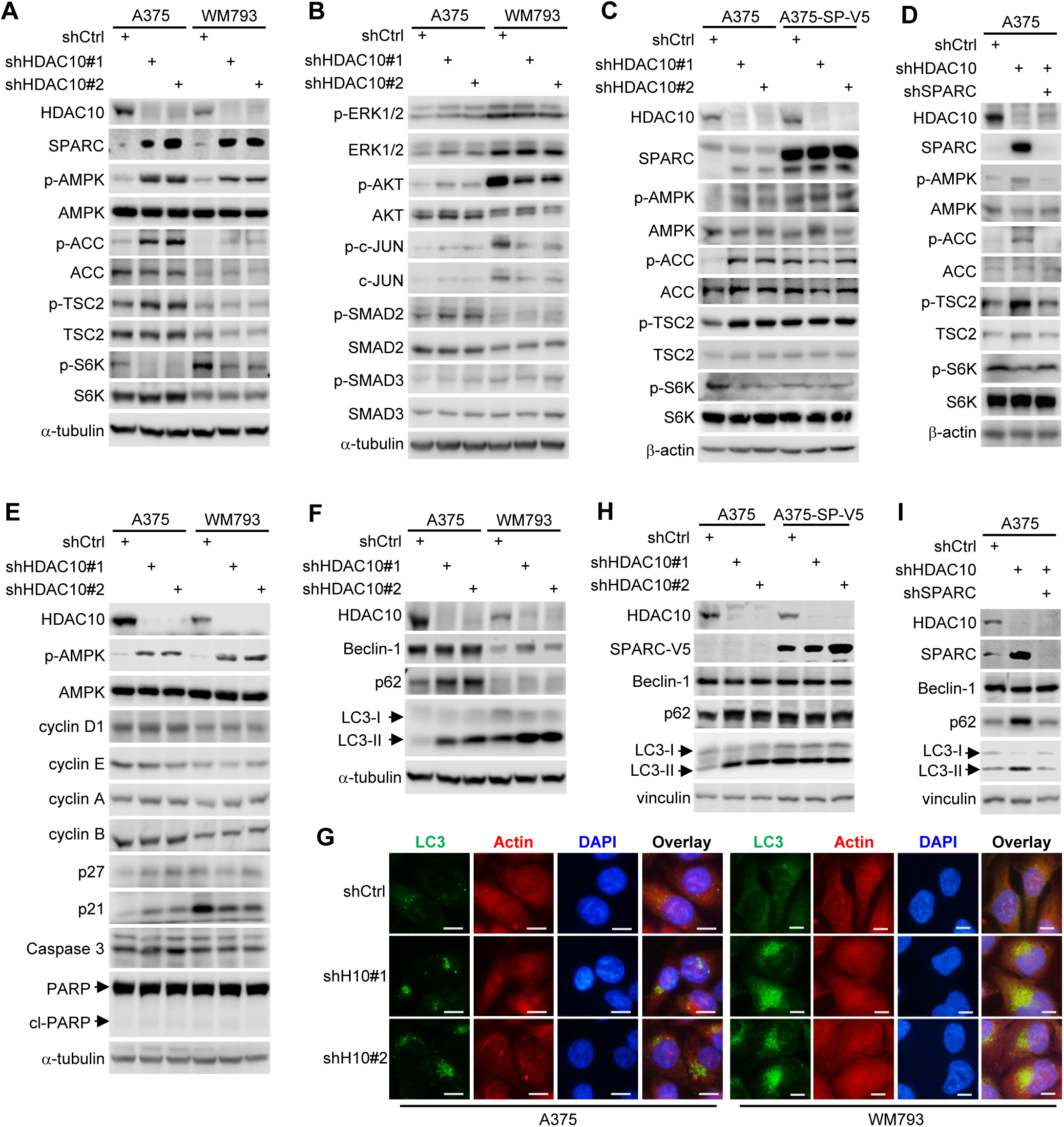
SPARC upregulation upon HDAC10 depletion induces AMPK activation and autophagic cell death. **(A)** A375 and WM793 cells were depleted of HDAC10. Cell lysates were immunoblotted with antibodies to indicated proteins. **(B)** Lysates from HDAC10-depleted and control A375 and WM793 cells were immunoblotted with antibodies to indicated proteins. **(C)** A375 cells were depleted of HDAC10 and/or overexpressed with SPARC. Cell lysates were immunoblotted with antibodies to indicated proteins. **(D)** Lysates from A375 cells depleted of HDAC10 and/or SPARC were immunoblotted with antibodies to indicated proteins. **(E)** Lysates from HDAC10-depleted or control A375 and WM793 cells were immunoblotted with antibodies to indicated proteins. **(F)** Lysates from HDAC10-depleted or control A375 and WM793 cells were immunoblotted with antibodies to Beclin-1, p62, and LC3. **(G)** HDAC10-depleted and control A375 and WM793 cells were stained for LC3 (green), actin (red), and nucleus (blue). **(H)** Lysates from A375 cells depleted of HDAC10 and/or overexpressed with V5-tagged SPARC were immunoblotted with antibodies to indicated proteins. **(I)** Lysates from A375 cells depleted of HDAC10 and/or SPARC were immunoblotted with antibodies to indicated proteins.

AMPK activation can inhibit cell growth and proliferation by inducing cell cycle arrest, apoptosis, and/or autophagy, but the outcomes may vary depending on the specific stimuli and cellular context (68–70). We examined the possible effects of the HDAC10-AMPK axis on the expression of key regulators for cell cycle progression (cyclins D1, E, A, and B, as well as p21 and p27) and apoptosis (PARP and Caspase 3). Surprisingly, although HDAC10 depletion led to AMPK activation, it did not significantly change the expression patterns of these regulators in either cell line, or consistently in both cell lines (Figure 5E). Thus, HDAC10 depletion caused cell growth inhibition does not result from cell cycle arrest or acute apoptosis, instead, is likely due to autophagic cell death. Supporting this hypothesis, HDAC10 depletion resulted in remarkable increases in the conversion of LC3-I to LC3-II and the expression of p62 (Figure 5F), indicating autophagy-like changes. In accordance with this, the LC3 autophagic puncta formation and accumulation in cells depleted of HDAC10 were enhanced (Figure 5G). Since SPARC is essential for AMPK activation upon HDAC10 depletion, we evaluated its impact on HDAC10 depletion-induced autophagy. The A375 cells depleted of HDAC10 were expressed with or depleted of SPARC and then were examined for the expression of LC3, p62, and Beclin-1. SPARC depletion largely attenuated the autophagy triggered by HDAC10 depletion (Figure 5H). In contrast, SPARC overexpression counteracted the effects of HDAC10 depletion and induced autophagy (Figure 5I). Together, these results indicate that the growth inhibition resulting from HDAC10 depletion is likely attributed to the activation of AMPK and subsequent autophagic cell death, via upregulation of SPARC.

### Expression of HDAC10 and SPARC is responsive to BRAF inhibition

Melanoma is the most aggressive form of skin cancer with a tendency to metastasize to other parts of the body if not detected and treated early. Approximately 50–60% of cutaneous melanomas harbor BRAF mutations, predominantly V600E. Targeted therapies that specifically inactivate mutated BRAF protein using BRAF inhibitor (BRAFi), such as vemurafenib (PLX4032), have yielded significant positive outcomes in patients with BRAF-mutated melanoma. Yet, the emergence of resistance to these drugs remains a major challenge in long-term treatment.

HDAC10 and SPARC are implicated in chemotherapy response across different types of cancer (10, 37, 71–74), but their roles in BRAFi resistance in melanoma are unexplored. Notably, the expression dynamics of both HDAC10 and SPARC are influenced by the BRAF activity state in melanoma cells carrying BRAF mutations. While BRAF activity minimally affects AMPK expression and activation, PLX4032 treatment, evidenced by decreased phospho-ERK1/2 levels, elevated HDAC10 at both protein and mRNA levels, subsequently reducing SPARC levels (Figures 6A, 6B). Thus, the BRAF activity state is crucial for the expression of HDAC10 and SPARC. Further, when exposed to PLX4032, HDAC10-depleted melanoma cells exhibited more pronounced inductions of both SPARC protein and mRNA (Figures 6C, 6D). These findings not only reiterate HDAC10’s suppressive impact on SPARC expression but also spotlight its potentially pivotal role in modulating SPARC response to PLX4032. However, since both A375 and WM793 cells display sensitivity to PLX4032, although HDAC10 knockdown appears to demonstrate an additional augmentation of sensitivity to the treatment (Figures 6E, 6F), it remains to be conclusively determined whether HDAC10 depletion intrinsically alters melanoma cell susceptibility to BRAFi.

**Figure 6.**
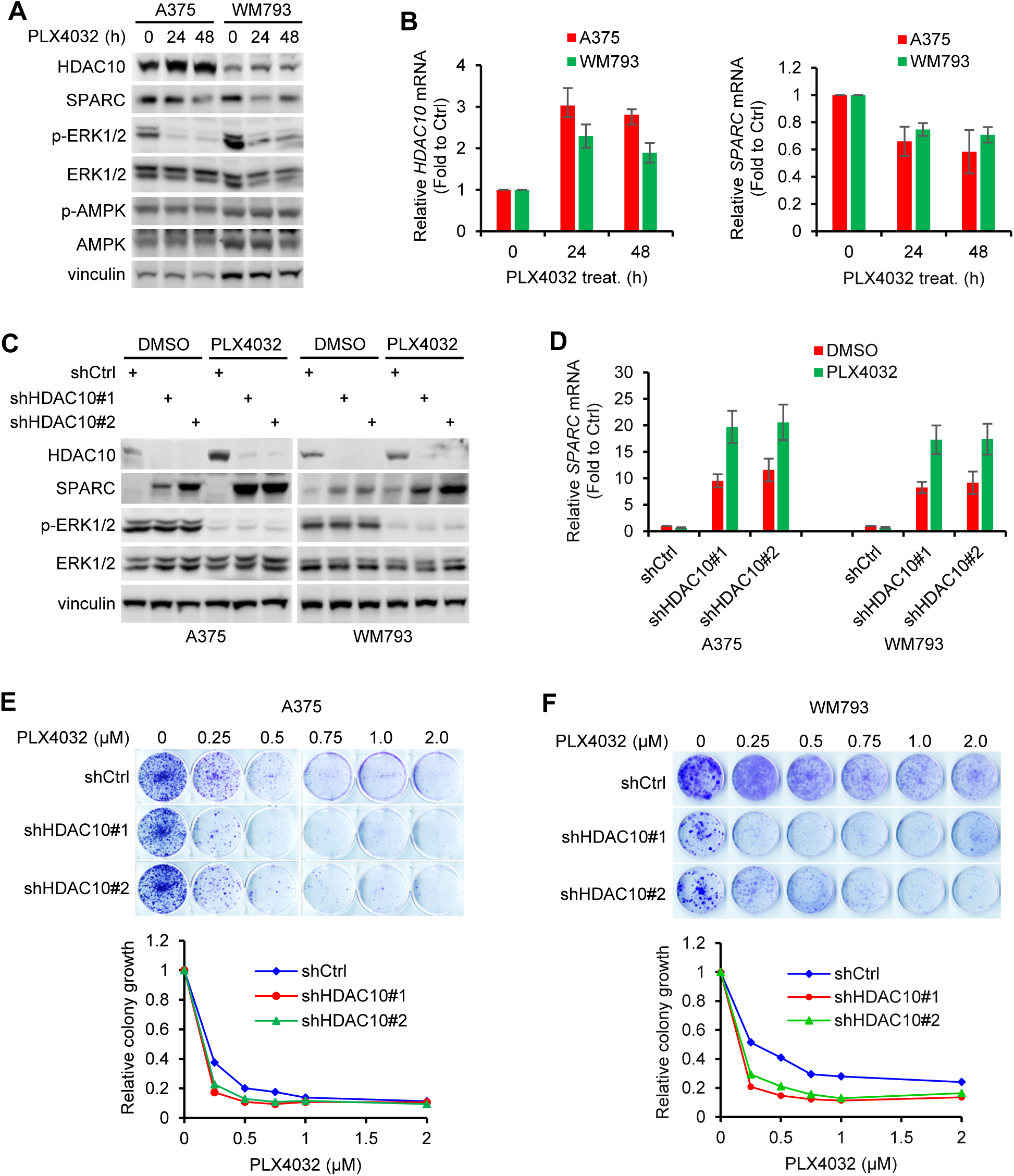
Expression of HDAC10 and SPARC is responsive to BRAF inhibition. **(A, B)** A375 and WM793 cells were treated with 1 µM PLX4032 for 24 h or 48 h. Cell lysates were subjected to immunoblotting with antibodies to indicated proteins (A). The relative mRNA levels of *HDAC10* and *SPARC* were quantified by qPCR (B). **(C, D)** HDAC10-depleted or control cells underwent PLX4032 treatment, then were immunoblotted with antibodies to indicated proteins (C). The relative *SPARC* mRNA were quantified by qPCR (D). **(E, F)** HDAC10-depleted and control cells were seeded on 6-well plates (2,000 cells for each well) and treated with indicated doses of PLX4032. Cells were stained with crystal violet (upper panel). Relative colony growth was quantified by Image J (lower panel).

### HDAC10 depletion overcomes BRAF inhibitor resistance partially through upregulating SPARC expression

We next developed a PLX4032-resistant cell line (A375-R) through chronic in vitro exposure to PLX4032. A375-R cells exhibited a markedly higher IC50 to PLX4032 compared to the parental A375 cells (A375-P) (Figure 7A). While A375-P cell proliferation was effectively inhibited by PLX4032, A375-R cells retained their proliferative vigor in its presence (Figures 7B, 7C).

**Figure 7.**
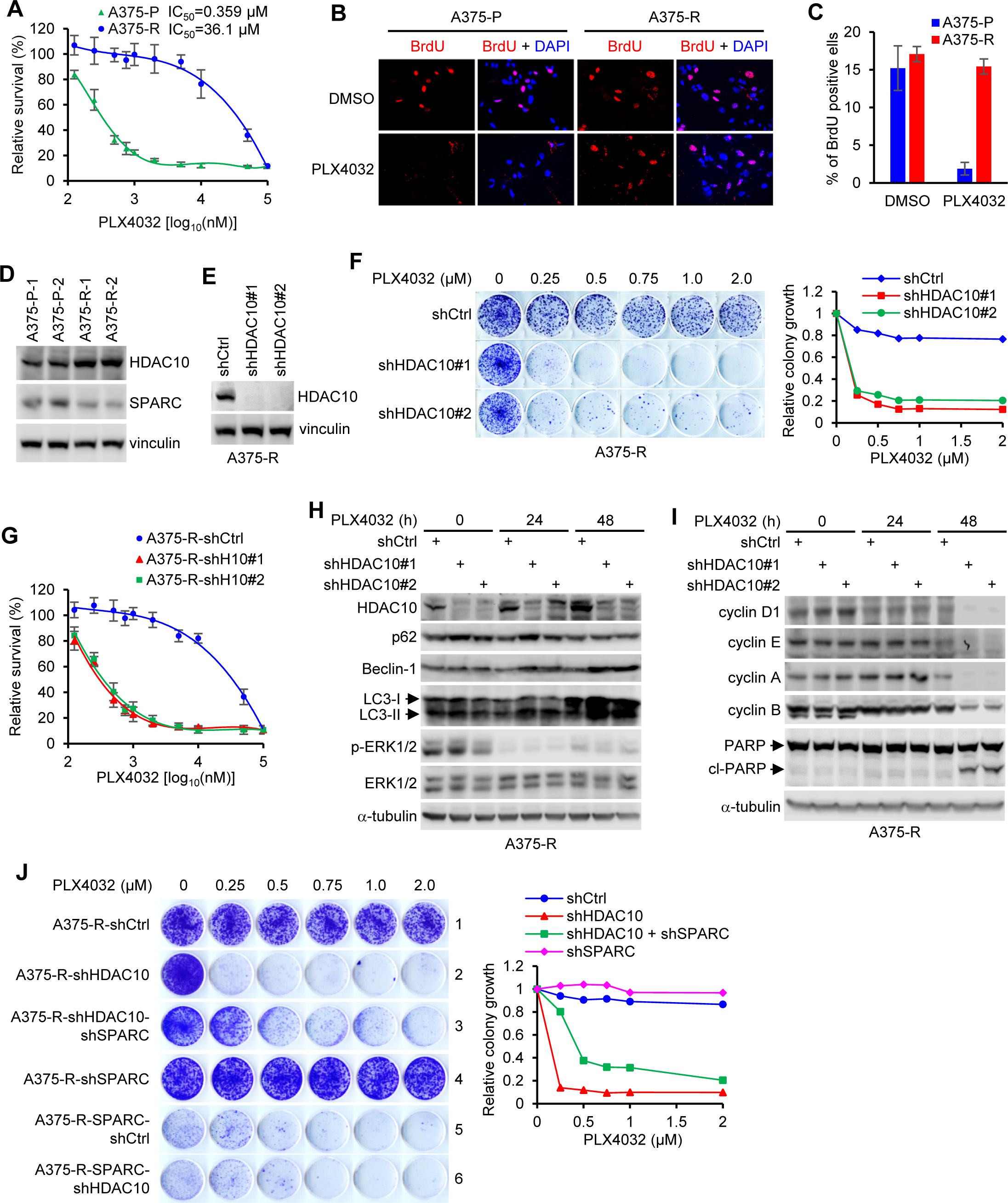
HDAC10 depletion overcomes BRAF inhibitor resistance partially through upregulating SPARC. **(A)** Parental (A375-P) and PLX4032-resistant (A375-R) A375 cells underwent gradient PLX4032 treatments for 72 h. Cell viability was ascertained using the CCK8 assay, with data displayed as mean ± SD over three repetitions. **(B, C)** Cells were treated with PLX4032 (2 µM) or the DMSO control for 24 h, and subjected to BrdU incorporation assay. Cells were stained for BrdU (red) and nucleus (blue). Representative BrdU staining images are provided (B), and BrdU incorporation rates are graphed as mean ± SD from three experiments (C). For each, at least 300 cells are counted. **(D)** Lysates from A375-P and A375-R cells were immunoblotted for HDAC10 and SPARC. **(E)** Lysates from HDAC10-depleted and control A375-R cells were immunoblotted for HDAC10 and vinculin. **(F)** HDAC10-depleted and control A375-R cells were seeded on 6-well plated, and treated with indicated doses of PLX4032 for 10 days. Cells were stained with crystal violet (left panel), and relative colony growth of cells was graphed (right panel). **(G)** HDAC10-depleted and control cells were treated with gradient doses of PLX4032 for 72 h. Relative cell viability was evaluated by CCK8 assay. **(H, I)** HDAC10-depleted and control A375-R cells were treated with PLX4032 for 24 h or 48 h. Cell lysates were immunoblotted with antibodies to indicated proteins. **(J)** A375-R cells were depleted of HDAC10, and/or were depleted of or overexpressed with SPARC. Cells were then treated with gradient doses of PLX4032 for 10 days. Cells were stained with crystal violet (upper panel) and relative colony growth of cells was graphed (lower panel).

Intriguingly, we observed an upregulation of HDAC10 and a concurrent downregulation of SPARC in A375-R cells in comparison to A375-P cells (Figure 7D). This finding, in conjunction with those presented in Figure 6, suggests that the shifts in the expression levels of HDAC10 and SPARC by BRAF inhibition are not transient or stochastic responses. Instead, they seem to exhibit relative stability, which could have significant and enduring implications for melanoma treatment strategies involving BRAFi. To gain insight into the potential influence of HDAC10 on BRAFi resistance, A375-R cells were depleted of HDAC10 (Figure 7E), subsequently exposed to various concentrations of PLX4032, and were evaluated for viability using colony formation and CCK8 assays. Depletion of HDAC10 resulted in a noteworthy reduction in colony growth (Figure 7F) and, consistently, a substantial restoration of sensitivity to PLX4032 (Figure 7G). In concordance with these findings, HDAC10 depletion led to an augmentation of cellular autophagy upon PLX4032 treatment, as evidenced by increased expression of key autophagic markers such as LC3, p62, and Beclin-1 (Figure 7H). Since the vulnerability to PLX4032 was observed in both colony formation and CCK8 assays, HDAC10 depletion may induce acute cell cycle arrest, potentially accompanied by cell death, in addition to observed autophagic response. Supporting this hypothesis, exposure to PLX4032 for 48 h significantly downregulated the expression of cyclins while upregulating the expression of cleaved PARP in HDAC10-depleted cells (Figure 7I). Thus, HDAC10 depletion may influence melanoma cell fate in diverse ways depending on the cellular context and the presence of other signaling pathways. In situations where cellular damage or stress is severe and the cell’s ability to adapt or recover is compromised without HDAC10, besides autophagy, acute cell cycle arrest and apoptosis may occur.

Finally, we asked if SPARC has a role in HDAC10-mediated regulation of PLX4032 resistance in A375-R cells. To this end, HDAC10-depleted or control A375-R cells were further depleted of or overexpressed with SPARC, then were evaluated for PLX4032 response. As expected, HDAC10 depletion rendered A375-R cells vulnerable to PLX4032 (Figure 7J, left panel, rows 1 and 2). Intriguingly, while the sole depletion of SPARC (SPARC was depleted in A375-R cells) had little effect on, the double depletion of SPARC (SPARC was depleted in HDAC10-depleted A375-R cells) resulted in a partial yet noticeable restoration of, sensitivity to the PLX4032 (Figure 7J, left panel, rows 3, 4). These results imply that HDAC10 likely involves PLX4032 response regulation through multiple pathways, besides SPARC signaling.

Additionally, these findings suggest that it is SPARC upregulation induced by HDAC10 depletion, but not the intrinsic SPARC expression, that plays a role in regulating PLX4032 response in A375-R cells. Supporting this notion, overexpression of SPARC markedly reduced colony growth and PLX4032 resistance in A375-R cells, regardless of the HDAC10 status (Figure 7J, left panel, rows 5, 6).

## DISCUSSION

HDAC10 belongs to the class IIb HDAC family, which was initially identified based on its sequence similarity to other class II HDACs (75–78). Particularly, it shares the highest structural resemblance with HDAC6, the other member of class IIb HDACs. Despite the similarity, they have distinct functions and subcellular localizations. HDAC6 is primarily cytoplasmic, mainly deacetylating cytosolic targets and guiding related processes (79, 80). HDAC10, on the other hand, is potentially more versatile with its presence both in the cytoplasm and the nucleus. This dual localization hints at a broader role for HDAC10, potentially in nuclear processes that HDAC6 might not partake in. HDAC10’s nuclear localization implies interactions with histones or transcriptional (co)factors, thereby influencing gene regulation. A case in point is that HDAC10 binds to the let-7f-2/miR-98 promoter region and causes histone H3 hypoacetylation, triggering a chain reaction involving the repression of let-7, upregulation of HMGA2, and ultimately, impacting cyclin A2 transcription (57). HDAC10’s versatility extends beyond histones. In melanocytes, HDAC10 forms complexes with Pax3 and KAP1, maintaining them in a deacetylated state, which influences the promoters of MITF, TRP-1, and TRP-2, ultimately driving melanogenesis (81). Furthermore, an intriguing aspect of HDAC10’s functionality is its ability to regulate gene expression without necessarily altering the acetylation states of histones or (co)factors. For example, in association with NCOR2, HDAC10 impacts DUB3 expression, thereby regulating BRD4 protein levels (82).

Our study delves deeper into HDAC10’s specific regulatory influence on SPARC expression; its depletion sharply elevates SPARC levels, while its overexpression does the opposite. The interplay between HDAC10 and histone modifications was another focal point of our research. We established a direct correlation between HDAC10 and H3K27ac, a modification often associated with gene activation, at *SPARC* regulatory elements. Harnessing the precision of a dCas9-mediated CRISPR approach, our study elucidated the competition between HDAC10 and p300/CBP, two crucial regulators of H3K27ac, at *SPARC* Enh2. These findings shed light on the intricate epigenetic mechanisms governing SPARC expression and underscore the potential of targeting HDAC10 in therapeutic strategies.

Although HDAC10 was identified over two decades ago (75–78, 83), the physiological function of HDAC10 as a lysine deacetylase remains controversial. In early studies, using either purified core histones or synthetic histone H4 peptide substrates, anti-FLAG immunoprecipitates derived from FLAG-tagged HDAC10 expressing cells exhibited histone deacetylase activities (75–78). Further, these same groups show that HDAC10 possesses transcriptional repression activity when targeted to promoters as a Gal4-fusion protein, though there were disagreements about whether repression by HDAC10 requires its intrinsic histone deacetylase activity. Later, using purified recombinant HDAC10, the enzymatic activity of HDAC10 could not be determined with a wide range of fluorophore-conjugated substrates (84, 85). These data raised the intriguing question of whether HDAC10 truly possesses lysine deacetylase activity or has a very limited number of substrates that may involve non-lysine and non-proteins. The answer came with the recent breakthrough discovery that HDAC10 is a robust polyamine deacetylase, with optimal catalytic activity and specificity for the hydrolysis of N(8)-acetylspermidine (86). While these recent results clearly demonstrate that recombinant HDAC10 has negligible lysine deacetylase activity in vitro, some reports support earlier findings that HDAC10 catalyzes protein deacetylations. For instance, HDAC10 catalyzes the deacetylation of non-histone proteins including the 70 kDa heat shock proteins (Hsc70/Hsp70) (81), RNA-binding protein EWS (EWSR1) (87), Yes-associated protein 1 (YAP1) (88), MutS homolog 2 (MSH2) (83), and single-stranded DNA binding protein 1 (hSSB1) (89). Conceivably, HDAC10 associates with other HDACs to indirectly deacetylate proteins in living cells, while purified HDAC10 uncoupled to other HDACs retains direct polyamine, but not lysine deacetylase activity. This possibility is reinforced by two early reports that HDAC10 interacts with HDAC2 and HDAC3 (75, 78). Also, HDAC10 has been shown to repress the promoter of CXCL10 by recruitment of EZH2 (90), OsHKT2 by interaction with OsPRR73 (91), DUB3 by complexing with NCOR2 (82), and MMP-2 and -9 by inhibiting RNA polymerase II (92).

Our observation that HDAC10 modulates the H3K27ac state at *SPARC* locus and represses its expression does not endorse that HDAC10 possesses intrinsic histone deacetylase activity. Rather, it fits well with the hypothesis that HDAC10 interacts with other HDACs or acetylation-modifying enzymes to indirectly regulate histone/protein modifications in cells. Alternatively, though non-mutually exclusive, with its robust polyamine deacetylase activity towards N(8)-acetylspermidine, it is possible that HDAC10 depletion leads to an increase in acetylspermidine levels, resulting in deregulated polyamine metabolism. This deregulation can potentially impact the activities of p300/CBP and other histone/protein acetyltransferases and, consequently, the acetylation state of histone and non-histone proteins, by affecting the availability of polyamine-derived acetyl groups for acetyltransferase enzymes (93, 94). Considering the widespread nature of (acetyl)spermidine, it is conceivable that HDAC10’s absence would influence the overall H3K27ac levels in cells. Yet, our work indicates histone deacetylation at specific genes, such as *SPARC*. We hypothesize that the observed rise in H3K27ac level at *SPARC* regulatory elements upon HDAC10 depletion is not due to spermidine’s deacetylation. Other alternative mechanisms could be at play: 1) HDAC10 may counteract or compete with p300/CBP and/or other HDACs at specific sites, thereby affecting the access of HATs and/or HDACs to the local histones surrounding the gene promoter or enhancers and influencing the acetylation state, and 2) HDAC10 could interact with transcription (co)factors, in turn, modulating the accessibility of p300 to histone H3. In addition to HDAC2 and SMRT, several such (co)factors, including TP53INP1 and SOX5, have been known to influence *SPARC* transcription and may work in tandem with HDAC10 (95–100). The precise mechanism by which HDAC10 influences the H3K27ac state at *SPARC* locus warrants further exploration.

SPARC is involved in various aspects of development and growth of melanoma. It exhibits both tumor-promoting and tumor-suppressing effects, depending on the context and stage of melanoma progression (10, 11). Our work indicates that the growth inhibition of melanoma cells is closely linked to SPARC’s upregulation following stable HDAC10 depletion. The transient depletion of HDAC10 may lead to severe mitotic catastrophe and cell cycle arrest in 2-D culture in some melanoma lines. After bypassing a proliferative crisis, the surviving HDAC10-depleted cells that restore a similar proliferative ability in 2-D monolayer culture retain an upregulation of SPARC. In this regard, the acute cell death and cell cycle arrest occurring at an early stage upon HDAC10 depletion is unlikely due to the upregulation of SPARC. In support of this perspective, the expression levels of cell cycle and apoptosis regulators remain relatively stable in these HDAC10-depleted cells, compared to that in the control cells. Instead, as time progresses post-HDAC10 depletion, the inhibitory effect on cell growth, attributed to the SPARC upsurge, seems to result from autophagic death.

Our study further deciphers the underlying relationship between HDAC10, SPARC, and AMPK; the HDAC10 depletion leads to the upregulation of SPARC, in turn, heightening the phosphorylation of AMPK. Corroboratively, overexpressing SPARC elevates AMPK phosphorylation, effectively overshadowing the influence of HDAC10 depletion. Inversely, SPARC depletion dampens the activation of AMPK upon HDAC10 depletion. In essence, the activation of AMPK signaling upon HDAC10 depletion hinges on the upregulation of SPARC.

SPARC’s multifaceted roles encompass aspects such as cell adhesion, migration, proliferation, and survival. In certain cell types, SPARC maintains an intracellular presence, implying it may have unexplored intracellular functions (22, 25, 101–105). SPARC’s roles in mediating AMPK activation, facilitating glucose metabolism, and interacting with metabolic pathways are well-documented (58, 59, 106). Moreover, SPARC-induced AMPK activation has been linked to autophagy (107). In our study, AMPK activation, triggered by HDAC10 depletion and concurrent SPARC upregulation, was found to potently induce autophagy, as reflected in the marked conversion of LC3-I to LC3-II. The HDAC10 depletion appears to negligibly impact cell apoptosis and cell cycle progression; the inhibition of melanoma cell growth is predominantly linked to autophagic cell death, a consequence of AMPK activation.

Our study also demonstrates that the expression levels of both HDAC10 and SPARC respond to the activity state of mutated BRAF in melanoma cells. Furthermore, our findings unveil a previously unrecognized role of the HDAC10-SPARC axis in modulating BRAFi response; HDAC10 depletion significantly re-sensitizes PLX4032-resistant melanoma cells to PLX4032 treatment, while SPARC depletion partially reverses this effect. For the first time, our findings underscore the significance of HDAC10 and SPARC in BRAFi resistance. These results also suggest that HDAC10 may influence BRAFi response through multiple downstream targets beyond SPARC. Further investigation is warranted to fully elucidate the underlying mechanisms.

In summary, our study marks a significant stride in understanding how HDAC10 controls SPARC expression, specifically through modulating H3K27ac state at key regulatory elements. Our work also establishes the HDAC10-SPARC axis as a critical regulator in melanoma cell growth and BRAFi resistance, offering insight into potential therapeutic targets for melanoma and related malignancies.

## ACKNOWLEDGEMENTS

We thank Cara Barmore for technical support and for critical reading of the manuscript. We also thank Ada Wong, Carol Ghezzi, Jameel Mughal, Jenna Clements, and TJ Laucevicius for helpful discussions and suggestions. This research was supported by NIH R01CA240529, R01AI153110, and R01CA270149 to E.S.

## AUTHOR CONTRIBUTIONS

H.L., Y.L., and E.S. conceived the project; H.L., Y.L., and C.P. performed the experiments and conducted the data analysis; S.Y. provided key reagents; H.L. and E.S. wrote the manuscript with significant input from all authors.

## DECLARATION OF GENERATIVE AI AND AI-ASSISTED TECHNOLOGIES IN THE WRITING PROCESS

During the final preparation of this work the authors used Chat Generative Pre-trained Transformer (ChatGPT) in order to improve the language and readability of this manuscript. After using this tool/service, the authors reviewed and edited the content as needed and take full responsibility for the content of the publication.

