## Supplemental Table and Figures for "HDAC10 blockade upregulates SPARC expression thereby repressing melanoma cell growth and BRAF inhibitor resistance"

**Supplemental Table S1. shRNA Target and Primer Sequences**

**<u>shRNA target sequences</u>**

| **shRNAs** | **Target sequences ( from 5' to 3' )** | **Sources** | **Identifier** |
|---|---|---|---|
| HDAC1-shRNA#1 | CGGTTAGGTTGCTTCAATCTA | Sigma | TRCN0000195467 |
| HDAC1-shRNA#2 | CCTAATGAGCTTCCATACAAT | Sigma | TRCN0000195103 |
| HDAC2-shRNA#1 | GCAAATACTATGCTGTCAATT | Sigma | TRCN0000004823 |
| HDAC2-shRNA#2 | CAGACTGATATGGCTGTTAAT | Sigma | TRCN0000195198 |
| HDAC3-shRNA#1 | CAAGAGTCTTAATGCCTTCAA | Sigma | TRCN0000194993 |
| HDAC3-shRNA#2 | CCTTCCACAAATACGGAAATT | Sigma | TRCN0000004825 |
| HDAC4-shRNA#1 | GCCAAAGATGACTTCCCTCTT | Sigma | TRCN0000004832 |
| HDAC4-shRNA#2 | GCCAAAGATGACTTCCCTCTT | Sigma | TRCN0000314665 |
| HDAC5-shRNA#1 | GCTAGAGAAAGTCATCGAGAT | Sigma | TRCN0000004835 |
| HDAC5-shRNA#2 | GCTCAAGAATGGATTTGCCAT | Sigma | TRCN0000004836 |
| HDAC6-shRNA#1 | CGGTAATGGAACTCAGCACAT | Sigma | TRCN0000004842 |
| HDAC6-shRNA#2 | CATCCCATCCTGAATATCCTT | Sigma | TRCN0000314976 |
| HDAC7-shRNA#1 | GATCCGGGTGCACAGTAAATA | Sigma | TRCN0000255687 |
| HDAC7-shRNA#2 | CAAGTAGTTGGAACCAGAGAA | Sigma | TRCN0000195442 |
| HDAC8-shRNA#1 | GCGTATTCTCTACGTGGATTT | Sigma | TRCN0000350469 |
| HDAC8-shRNA#2 | GCGTATTCTCTACGTGGATTT | Sigma | TRCN0000004851 |
| HDAC9-shRNA#1 | CAAACTGCTTTCGAAATCTAT | Sigma | TRCN0000195059 |
| HDAC9-shRNA#2 | GAGCAGTTAATAGGCTTTAAA | Sigma | TRCN0000196384 |
| HDAC10-shRNA#1 | CCTGTACCTCTTAGATGGGAT | Sigma | TRCN0000004860 |
| HDAC10-shRNA#2 | GTGTTCAACAACGTGGCCATA | Sigma | TRCN0000004861 |
| HDAC11-shRNA#1 | CCCGACGTGGTGGTATACAAT | Sigma | TRCN0000199149 |
| HDAC11-shRNA#2 | GCGCTATCTTAATGAGCTCAA | Sigma | TRCN0000330863 |
| SPARC-shRNA | CGGTTGTTCTTTCCTCACATT | Sigma | TRCN0000008709 |
| p300-shRNA | CAATTCCGAGACATCTTGAGA | Sigma | TRCN0000009883 |
| BRD4-shRNA | CCTGGAGATGACATAGTCTTA | Sigma | TRCN0000021427 |
| Control-shRNA | GCAAGCTGACCCTGAAGTTCAT | Addgene | #30323 |

**<u>Primer sequences for RT-qPCR</u>**

| **Primers** | **Sequences ( from 5' to 3' )** | **Sources** | **Identifier** |
|---|---|---|---|
| HDAC1-RT-F | CGAATCCGCATGACTCATAA | IDT | This paper |
| HDAC1-RT-R | CATCTCCTCAGCATTGGCTT | IDT | This paper |
| HDAC2-RT-F | ATGGCGTACAGTCAAGGAGG | IDT | This paper |
| HDAC2-RT-R | ATGAGGCTTCATGGGATGAC | IDT | This paper |
| HDAC3-RT-F | GCAAGGCTTCACCAAGAGTC | IDT | This paper |
| HDAC3-RT-R | CTGTGTAACGCGAGCAGAAC | IDT | This paper |
| HDAC4-RT-F | CGTGGAAATTTTGAGCCATT | IDT | This paper |
| HDAC4-RT-R | CTGGTCTCGGCCAGAAAGT | IDT | This paper |
| HDAC5-RT-F | GGGAACCATCCTTGGAAATC | IDT | This paper |
| HDAC5-RT-R | GAACTGGGCATGGCTCTTG | IDT | This paper |
| HDAC6-RT-F | CCGGAGGGTCCTTATCGTAG | IDT | This paper |
| HDAC6-RT-R | GCGGTGGATGGAGAAATAGA | IDT | This paper |
| HDAC7-RT-F | CTGCATTGGAGGAATGAAGCT | IDT | This paper |
| HDAC7-RT-R | CTGGCACAGCGGATGTTTG | IDT | This paper |
| HDAC8-RT-F | ATACTTGACCGGGGTCATCC | IDT | This paper |
| HDAC8-RT-R | GCGTGATTTCCAGCACATAA | IDT | This paper |
| HDAC9-RT-F | GTACAGAAGTAAAGCAGAAG | IDT | This paper |
| HDAC9-RT-R | AGAGCTTTGATCCAATGATG | IDT | This paper |

| | | | |
|---|---|---|---|
| HDAC10-RT-F | CTCGGCTTCACTGTCAACCT | IDT | This paper |
| HDAC10-RT-R | GTCAAATCCTGCCGAGACCA | IDT | This paper |
| HDAC11-RT-F | GCACACGAGGCGCTATCTTA | IDT | This paper |
| HDAC11-RT-R | AAGGAAGTTGGGGAGGAAGA | IDT | This paper |
| SPARC-RT-F | ACATAAGCCCAGTTCATCACCA | IDT | This paper |
| SPARC-RT-R | ACAACCGATTCACCAACTCCA | IDT | This paper |
| GAPDH-RT-F | AAGGTGAAGGTCGGAGTCAA | IDT | This paper |
| GAPDH-RT-R | AATGAAGGGGTCATTGATGG | IDT | This paper |

**Primer sequences for generation of plentiCRISPR-gRNA targeting *HDAC10***

| **Primers** | **Sequences ( from 5' to 3' )** | **Sources** | **Identifier** |
|---|---|---|---|
| HDAC10-gRNA-F | CACCGGACGCTCGATCTCGCACTCG | IDT | This paper |
| HDAC10-gRNA-R | AAACCGAGTGCGAGATCGAGCGTCC | IDT | This paper |

**Primers sequences for generation of dCas9/CRISPR-gRNAs targeting *SPARC loci***

| **Primers** | **Sequences (from 5' to 3')** | **Sources** | **Identifier** |
|---|---|---|---|
| Pro-gRNA1-F | CACCGGATCTGCCCTGGGCTGACCA | IDT | This paper |
| Pro-gRNA1-R | AAACTGGTCAGCCCAGGGCAGATCC | IDT | This paper |
| Pro-gRNA2-F | CACCGGTCTGTCCCTTGGTCAGCCC | IDT | This paper |
| Pro-gRNA2-R | AAACGGGCTGACCAAGGGACAGACC | IDT | This paper |
| Pro-gRNA3-F | CACCGGAATGATGCAGTCTGTCCCT | IDT | This paper |
| Pro-gRNA3-R | AAACAGGGACAGACTGCATCATTCC | IDT | This paper |
| Pro-gRNA4-F | CACCGGACTGCATCATTCAGTCCAC | IDT | This paper |
| Pro-gRNA4-R | AAACGTGGACTGAATGATGCAGTCC | IDT | This paper |
| Pro-gRNA5-F | CACCGGAGATGTAACATTTGCCCTG | IDT | This paper |
| Pro-gRNA5-R | AAACCAGGGCAAATGTTACATCTCC | IDT | This paper |
| Enh1-gRNA1-F | CACCGGCTCAGTCTGGTGCTGATAA | IDT | This paper |
| Enh1-gRNA1-R | AAACTTATCAGCACCAGACTGAGCC | IDT | This paper |
| Enh1-gRNA2-F | CACCGAAGGATATAGCTGCTCAGTC | IDT | This paper |
| Enh1-gRNA2-R | AAACGACTGAGCAGCTATATCCTTC | IDT | This paper |
| Enh1-gRNA3-F | CACCGCAGTCTGGTGCTGATAATGG | IDT | This paper |
| Enh1-gRNA3-R | AAACCCATTATCAGCACCAGACTGC | IDT | This paper |
| Enh1-gRNA4-F | CACCGTAATGGTGGTAGTGATCAAA | IDT | This paper |
| Enh1-gRNA4-R | AAACTTTGATCACTACCACCATTAC | IDT | This paper |
| Enh1-gRNA5-F | CACCGCTATATCCTTTTATTAATCA | IDT | This paper |
| Enh1-gRNA5-R | AAACTGATTAATAAAAGGATATAGC | IDT | This paper |
| Enh2-gRNA1-F | CACCGGCTCCACACCCAGCCTTGGG | IDT | This paper |
| Enh2-gRNA1-R | AAACCCCAAGGCTGGGTGTGGAGCC | IDT | This paper |
| Enh2-gRNA2-F | CACCGGAGGAAAGGATATTTAAAGC | IDT | This paper |
| Enh2-gRNA2-R | AAACGCTTTAAATATCCTTTCCTCC | IDT | This paper |
| Enh2-gRNA3-F | CACCGGCAGAGCTAGGGGTAGACCT | IDT | This paper |
| Enh2-gRNA3-R | AAACAGGTCTACCCCTAGCTCTGCC | IDT | This paper |
| Enh2-gRNA4-F | CACCGGGCAGAGCTAGGGGTAGACC | IDT | This paper |
| Enh2-gRNA4-R | AAACGGTCTACCCCTAGCTCTGCCC | IDT | This paper |
| Enh2-gRNA5-F | CACCGGCAGTCAGTGGCAGAGCTAG | IDT | This paper |
| Enh2-gRNA5-R | AAACCTAGCTCTGCCACTGACTGCC | IDT | This paper |
| Enh3-gRNA1-F | CACCGGAGATCGACTAGGACTCCAT | IDT | This paper |
| Enh3-gRNA1-R | AAACATGGAGTCCTAGTCGATCTCC | IDT | This paper |
| Enh3-gRNA2-F | CACCGATGTCGCTGTACTAAACCTT | IDT | This paper |
| Enh3-gRNA2-R | AAACAAGGTTTAGTACAGCGACATC | IDT | This paper |

| | | | |
|---|---|---|---|
| Enh3-gRNA3-F | CACCGTACAGTTGTACTCACACCTA | IDT | This paper |
| Enh3-gRNA3-R | AAACTAGGTGTGAGTACAACTGTAC | IDT | This paper |
| Enh3-gRNA4-F | CACCGACAACTGTATGCTGAGCCAA | IDT | This paper |
| Enh3-gRNA4-R | AAACTTGGCTCAGCATACAGTTGTC | IDT | This paper |

**Primer sequences for chromatin immunoprecipitation (ChIP)-qPCR**

| **Primers** | **Sequences ( from 5' to 3' )** | **Sources** | **Identifier** |
|---|---|---|---|
| Control-ChIP-F | ATGTACTGGGGGTGTACGGA | IDT | This paper |
| Control-ChIP-R | CCAGCCACAGCCTTAGAACT | IDT | This paper |
| Pro-ChIP-F | TGAGTCGGTTTAGGCAGCAG | IDT | This paper |
| Pro-ChIP-R | TCCTCCTCTTTTTGCGGTGG | IDT | This paper |
| Enh1-ChIP-F | CCCAAGACTAGTGTCCAGCC | IDT | This paper |
| Enh1-ChIP-R | CGGCGAAGAAAGGTAGTGGA | IDT | This paper |
| Enh2-ChIP-F | CACCCACTGGCTCAGCTAAA | IDT | This paper |
| Enh2-ChIP-R | CAGCCTGTAATAGGCGAGGG | IDT | This paper |
| Enh3-ChIP-F | AGATGGGGGAACCAAGGTCT | IDT | This paper |
| Enh3-ChIP-R | GCCAGGGGGTCTTGTTCTAC | IDT | This paper |

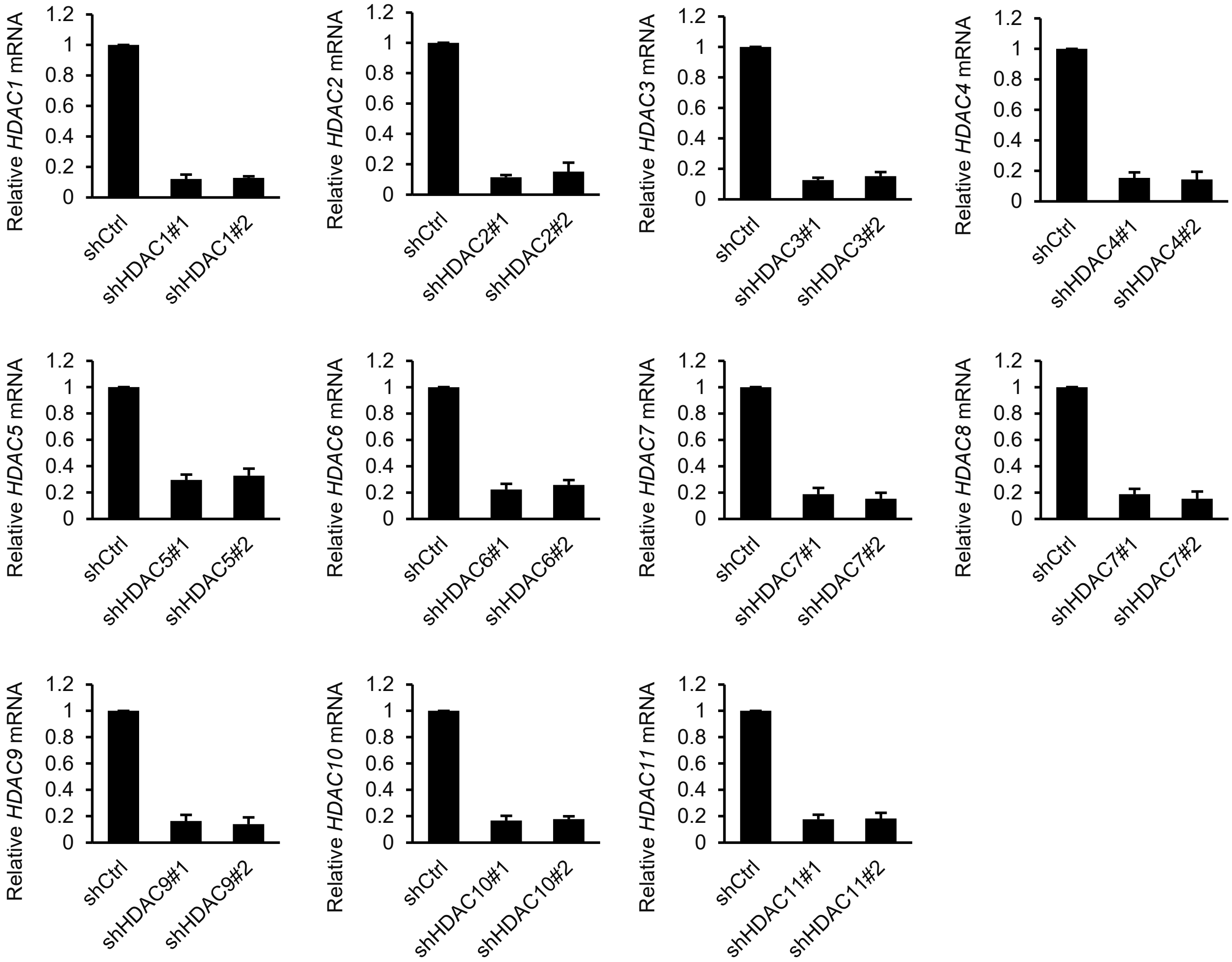


**Supplemental Figure S1. Evaluation of the knockdown of HDAC1–11 in A375 melanoma cells by qPCR.**

A375 melanoma cells were infected with lentiviruses harboring shRNA targeting individual HDAC member (HDACs 1–11) and control. Relative mRNA level of each HDAC was determined by qPCR.

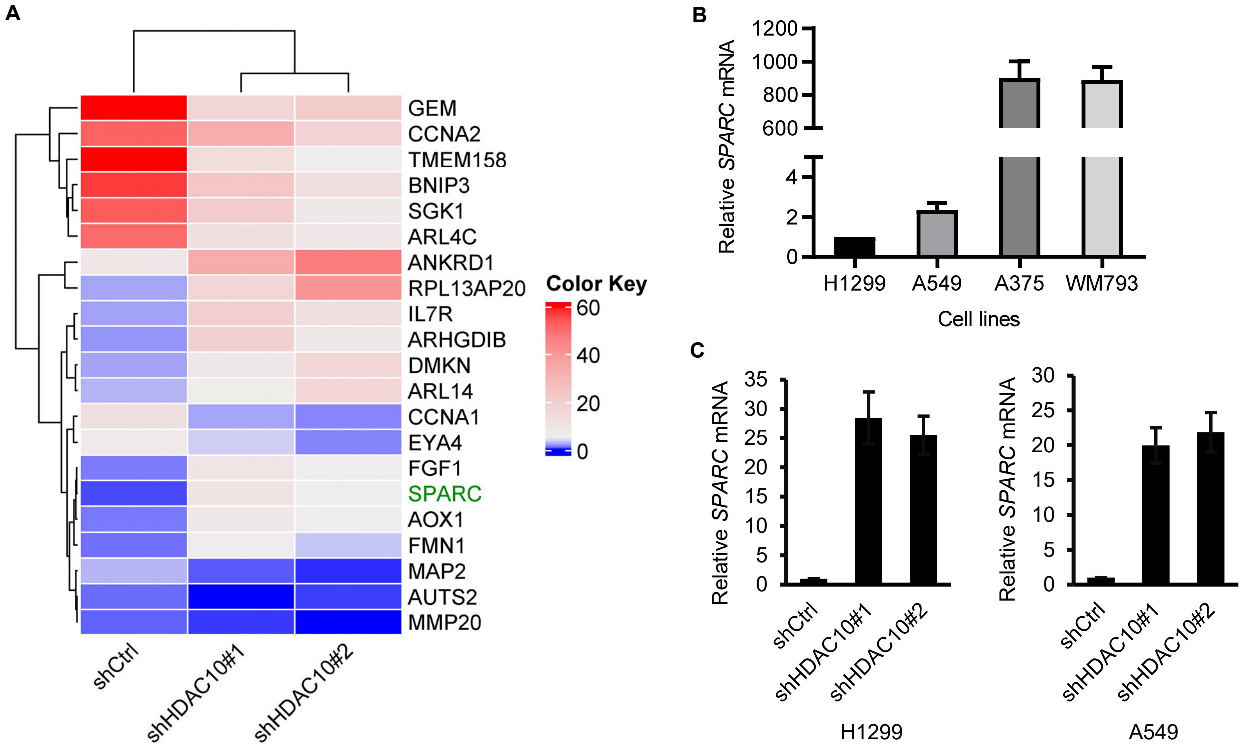


**Supplemental Figure S2. HDAC10 depletion upregulates *SPARC* mRNA level in lung cancer cells.**

(A) The top differential expression genes upon HDAC10 depletion in H1299 lung cancer cells by RNA-seq.

(B) Comparison of relative mRNA levels of *SPARC* between lung cancer (H1299 and A549) and melanoma (A375 and WM793) cell lines.

(C) HDAC10 depletion upregulates the mRNA level of *SPARC* in lung cancer cell lines.
